# Emergence of resistance to succinate dehydrogenase inhibitor fungicides in *Pyrenophora teres* f. *teres* and *P. teres* f. *maculata* in Australia

**DOI:** 10.1101/2023.04.23.537974

**Authors:** W.J. Mair, H. Wallwork, T.A. Garrard, J. Haywood, N. Sharma, K.N. Dodhia, R.P. Oliver, F.J. Lopez-Ruiz

## Abstract

The net blotches are among the most economically significant diseases of barley worldwide. There are two forms of the disease: net-form net-blotch (NFNB, causal agent *Pyrenophora teres* f. sp. *teres* [*Ptt*]) and spot-form net blotch (SFNB, causal agent *Pyrenophora teres* f. sp. *maculata* [*Ptm*]). Alongside varietal choice and cultural practices, fungicides form an important part of the regime for net blotch control. The succinate dehydrogenase inhibitors (SDHIs) are a key class of fungicides used in net blotch management. However, resistance to this group of compounds has emerged in the net blotches in recent years. Here, we describe the first cases of resistance to SDHIs in Australian populations of net blotches. This study was prompted by reports of field failures of SDHI fungicides in controlling NFNB in South Australia and SFNB in Western Australia. Target site mutations in the *Sdh* complex genes, previously associated with reduced sensitivity in European net blotch populations, were found in Australian isolates, and two mutations which have not been previously observed in *P. teres*, are also described. The mutations found in *Ptt* included H134R and S135R in *SdhC*; and H134Y and D145G in *SdhD*; the *SdhC*-H134R mutation was the most frequently observed. In *Ptm*, the mutations found included H277L in *SdhB*; S73P, N75S, H134R and S135R in *SdhC*; and D145G in *SdhD*; the *SdhC*-N75S mutation was the most common. These mutations were correlated with reduced *in vitro* SDHI fungicide sensitivity by microtiter assay. The highest resistance factors to fluxapyroxad and bixafen, the most important SDHI fungicides for net blotch control in Australia, were associated with the *SdhC*-H134R and *SdhC*-S135R mutations in *Ptt*, and with the *SdhB*-H277L, *SdhC*-H134R, and *SdhC*-S135R mutations in *Ptm*. Modelling of the *P. teres* Sdh complex showed that the two novel mutations, H277L in *SdhB* and H134Y in *SdhD*, result in a highly altered binding mode and lower binding affinity of the SDHI compound compared to the wild-type.

## 1. Introduction

The net blotch diseases are among the most economically significant diseases of barley (*Hordeum vulgare* L.) worldwide (Akhavan *et al*., 2017). There are two forms of the disease: net-form net-blotch (NFNB, causal agent *Pyrenophora teres* f. sp. *teres* (*Ptt*), Drechsler, anamorph *Drechslera teres* [Sacc.] Shoemaker) and spot-form net blotch (SFNB, causal agent *Pyrenophora teres* f. sp. *maculata* (*Ptm*), Smedegaard-Petersen) (Liu *et al*., 2011). The two forms are distinguished by their symptoms on host barley, with NFNB developing a distinct network pattern of dark brown striations, while SFNB produces circular to elliptical dark brown spots surrounded by a zone of chlorosis (Liu *et al*., 2011). The two *formae speciales* are closely related but genetically distinct and are considered as different species (Ellwood *et al*., 2012). In Australia, the net blotches are the most important diseases of barley with a combined losses estimated at AUD$62 million per annum, and with potential losses in the absence of current control measures estimated at AUD$309 million per annum (Murray & Brennan, 2010). Along with varietal choice and cultural practices, fungicides form an important part of the regime for net blotch control (Murray & Brennan, 2010; Rehfus *et al*., 2016; Akhavan *et al*., 2017).

Several cases of resistance to fungicides have been reported previously for the net blotch diseases. In Australia, laboratory and field resistance to some fungicides within the demethylase inhibitor (DMI) group has been reported in the net-form since 2013 (Mair *et al*., 2016a) and the spot-form since 2016 (Mair *et al*., 2020). In both of the forms, resistance was associated with the point mutation F489L in the DMI target enzyme *Cyp51A* as well as with overexpression of the target gene. There have been other reports of DMI resistance in *P. teres* from New Zealand (Sheridan *et al*., 1985), the United Kingdom (Sheridan *et al*., 1987), and Finland (Serenius & Manninen, 2008), although the molecular mechanisms were not described in these cases. Partial resistance to the Quinone-outside inhibitor (QoI) class has been reported in *P. teres* in Europe since 2003, associated with the point mutation F129L in the QoI target cytochrome *b* (Sierotzki *et al*., 2007). Since 2012, resistance has been observed in European *P. teres* populations towards the succinate dehydrogenase inhibitor (SDHI) class of fungicides (Rehfus *et al*., 2016).

The SDHIs (carboxamides, FRAC Code C2 or Group 7) are a structurally diverse class of compounds, all having in common an amide bond (Sierotzki & Scalliet, 2013b). These compounds act by targeting the enzyme complex succinate dehydrogenase (SDH, respiratory complex II or succinate-ubiquinone reductase), which couples the oxidation of succinate to fumarate with the reduction of ubiquinone to ubiquinol (Stammler *et al*., 2015). This enzyme complex therefore plays a key role in both the mitochondrial electron transport chain and the tricarboxylic acid (Krebs) cycle (Sierotzki & Scalliet, 2013b). The SDH enzyme complex consists of four subunits: subunit A (SdhA) is a flavoprotein that catalyses succinate to fumarate oxidation, subunit B (SdhB) is an iron-sulphur protein with three iron-sulphur clusters involved in electron transport, subunits C (SdhC) and D (SdhD) are complexed with a haem b group and form a membrane-anchoring domain (Stammler *et al*., 2015). The reduction of ubiquinone to ubiquinol occurs in the hydrophobic, highly-conserved ubiquinone-binding pocket (Q-site), formed from the B, C and D subunits (Stammler *et al*., 2015). The SDHIs bind deeper within the Q-site than does ubiquinone (Stammler *et al*., 2015), and interrupt the transfer of electrons from the iron-sulphur clusters to the substrate (Sierotzki & Scalliet, 2013b).

Amino acid substitutions affecting the binding affinity of SDHI fungicides have been found within the B, C and D subunits that together form the ubiquinone-binding pocket, although the mutated residues are not necessarily proximal to the actual Q-site (Stammler *et al*., 2015). Multiple mutations have been found occurring together, either within the same subunit (Pearce *et al*., 2019), or across different subunits (Rehfus, 2018). Non-target site resistance has also been reported (Yamashita & Fraaije, 2018), as well as reduced sensitivity from the overexpression of efflux pumps (Kretschmer *et al*., 2009; Omrane *et al*., 2015; Sang *et al*., 2015; Samaras *et al*., 2020). In *Zymoseptoria tritici*, resistance has also been associated with additional paralogs of subunit C (Steinhauer *et al*., 2019). In *P. teres*, the first detections of SDHI resistance in European populations were linked to the mutation H277Y in the SDH subunit B gene (Rehfus *et al*., 2016). Resistance increased substantially in frequency in subsequent years, and later isolates carried one of a number of mutations including *SdhB*-H277Y, *SdhC*-N75S, *SdhC*-H134R, *SdhC*-S135R, *SdhD*-H134R, and *SdhD*-D145G (Rehfus *et al*., 2016).

SDHIs have been used extensively in crop protection since the 1960s (Zhang *et al*., 2015). However, the first-generation SDHIs, such as carboxin, are effective mainly against Basidiomycetes and have limited activity towards other fungi (Avenot & Michailides, 2010). In contrast, the second-generation SDHIs have a broader spectrum of activity, including the Ascomycetes (Avenot & Michailides, 2010). Boscalid, the first of the second-generation SDHIs (Sierotzki & Scalliet, 2013a), was introduced in Australia in 2003, mainly for control of horticultural and viticultural diseases (APVMA, 2022). Fluxapyroxad was the first of the second-generation SDHIs for use on cereals in Australia, registered as a seed treatment in 2015 and introduced the following year (Platz *et al*., 2017). Bixafen, the first foliar SDHI for use on cereals, followed in 2018 (Bayer, 2018).

This study was prompted by reports of field failures of fluxapyroxad in controlling net-form and spot-form net blotches in South Australia (SA) and Western Australia (WA), respectively. The objectives of this study was to assess the *in vitro* sensitivity of *Ptt* and *Ptm* isolates towards SDHI fungicides, and to correlate reductions in sensitivity with mutations in the target *Sdh* genes. A molecular method for distinguishing the two *formae speciales* is also presented.

## 2. Materials and Methods

### 2.1 Fungal isolates

Barley samples from 2015-2020 were sourced from a combination of field trips and a network of collaborators. Reisolation of monoconidial *P. teres* strains was performed as previously described (Mair *et al*., 2020). Historical isolates from 1996 to 2012 were provided by Simon Ellwood (Centre for Crop and Disease Management, Curtin University). Isolates included in this study are listed in Table 1.

**Table 1.**
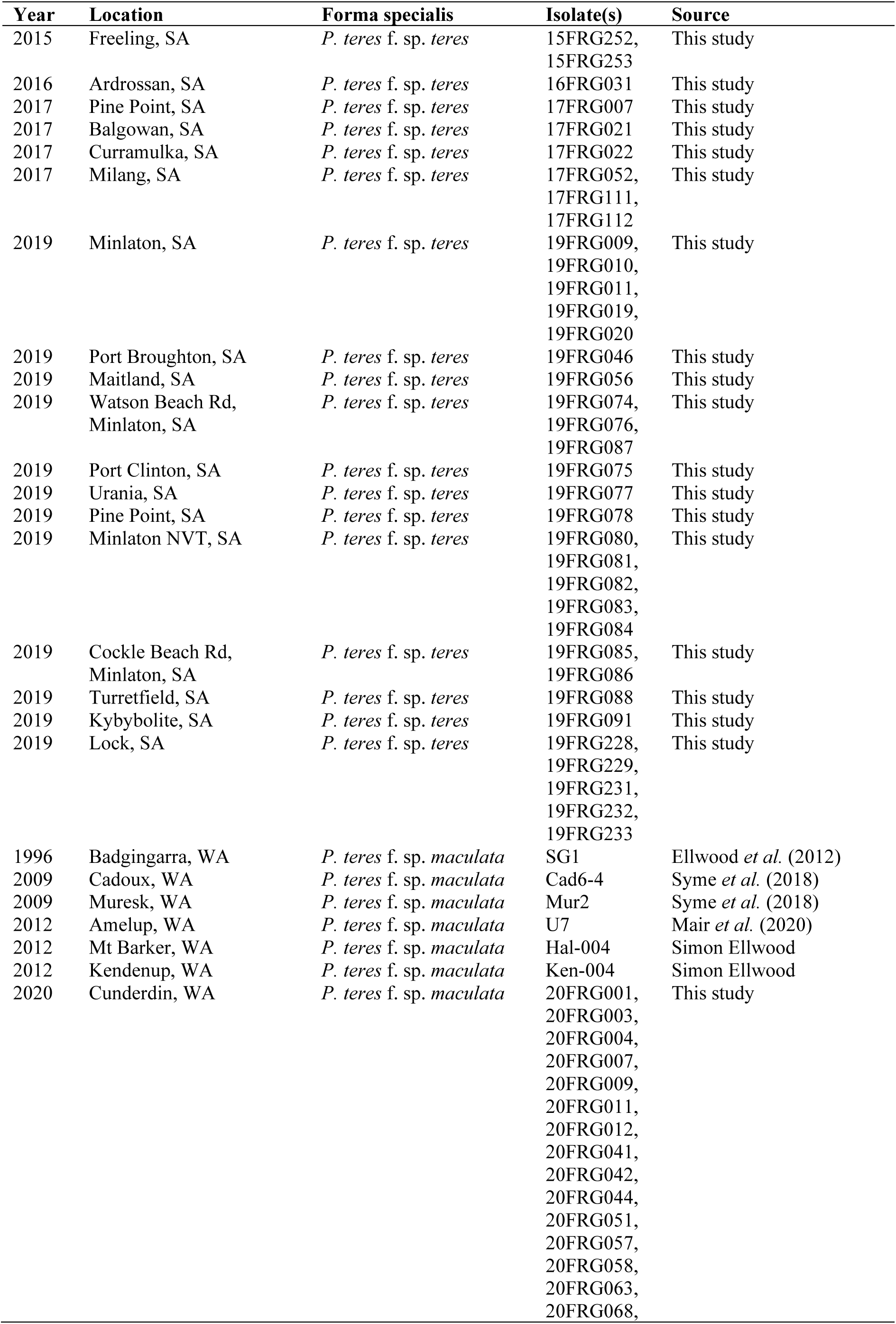
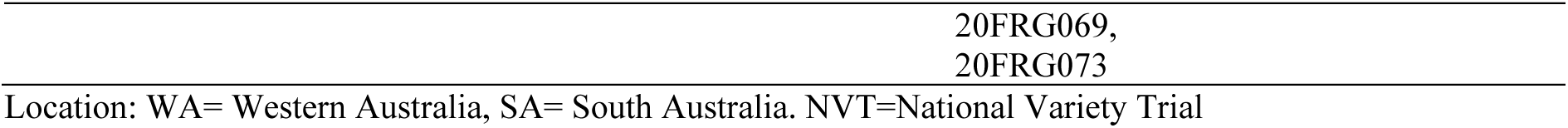
Isolates described in this study.

| Year | Location | Forma specialis | Isolate(s) | Source |
| --- | --- | --- | --- | --- |
| 2015 | Freeling, SA | *P. teres* f. sp. *teres* | 15FRG252,<br>15FRG253 | This study |
| 2016 | Ardrossan, SA | *P. teres* f. sp. *teres* | 16FRG031 | This study |
| 2017 | Pine Point, SA | *P. teres* f. sp. *teres* | 17FRG007 | This study |
| 2017 | Balgowan, SA | *P. teres* f. sp. *teres* | 17FRG021 | This study |
| 2017 | Curramulka, SA | *P. teres* f. sp. *teres* | 17FRG022 | This study |
| 2017 | Milang, SA | *P. teres* f. sp. *teres* | 17FRG052,<br>17FRG111,<br>17FRG112 | This study |
| 2019 | Minlaton, SA | *P. teres* f. sp. *teres* | 19FRG009,<br>19FRG010,<br>19FRG011,<br>19FRG019,<br>19FRG020 | This study |
| 2019 | Port Broughton, SA | *P. teres* f. sp. *teres* | 19FRG046 | This study |
| 2019 | Maitland, SA | *P. teres* f. sp. *teres* | 19FRG056 | This study |
| 2019 | Watson Beach Rd,<br>Minlaton, SA | *P. teres* f. sp. *teres* | 19FRG074,<br>19FRG076,<br>19FRG087 | This study |
| 2019 | Port Clinton, SA | *P. teres* f. sp. *teres* | 19FRG075 | This study |
| 2019 | Urania, SA | *P. teres* f. sp. *teres* | 19FRG077 | This study |
| 2019 | Pine Point, SA | *P. teres* f. sp. *teres* | 19FRG078 | This study |
| 2019 | Minlaton NVT, SA | *P. teres* f. sp. *teres* | 19FRG080,<br>19FRG081,<br>19FRG082,<br>19FRG083,<br>19FRG084 | This study |
| 2019 | Cockle Beach Rd,<br>Minlaton, SA | *P. teres* f. sp. *teres* | 19FRG085,<br>19FRG086 | This study |
| 2019 | Turretfield, SA | *P. teres* f. sp. *teres* | 19FRG088 | This study |
| 2019 | Kybybolite, SA | *P. teres* f. sp. *teres* | 19FRG091 | This study |
| 2019 | Lock, SA | *P. teres* f. sp. *teres* | 19FRG228,<br>19FRG229,<br>19FRG231,<br>19FRG232,<br>19FRG233 | This study |
| 1996 | Badgingarra, WA | *P. teres* f. sp. *maculata* | SG1 | Ellwood *et al.* (2012) |
| 2009 | Cadoux, WA | *P. teres* f. sp. *maculata* | Cad6-4 | Syme *et al.* (2018) |
| 2009 | Muresk, WA | *P. teres* f. sp. *maculata* | Mur2 | Syme *et al.* (2018) |
| 2012 | Amelup, WA | *P. teres* f. sp. *maculata* | U7 | Mair *et al.* (2020) |
| 2012 | Mt Barker, WA | *P. teres* f. sp. *maculata* | Hal-004 | Simon Ellwood |
| 2012 | Kendenup, WA | *P. teres* f. sp. *maculata* | Ken-004 | Simon Ellwood |
| 2020 | Cunderdin, WA | *P. teres* f. sp. *maculata* | 20FRG001,<br>20FRG003,<br>20FRG004,<br>20FRG007,<br>20FRG009,<br>20FRG011,<br>20FRG012,<br>20FRG041,<br>20FRG042,<br>20FRG044,<br>20FRG051,<br>20FRG057,<br>20FRG058,<br>20FRG063,<br>20FRG068, | This study |

Location: WA= Western Australia, SA= South Australia. NVT=National Variety Trial

### 2.2 Species identification

Two transcribed genetic regions, each unique to either *Ptt* (GenBank accession …) or *Ptm* (GenBank accession KX909560.1) were provided by Dr Simon Ellwood (pers. communication). A pair of primers per sequence were designed using PrimerExplorer v.4 (Eiken, Japan) with default settings and ordered from Macrogen (Seoul, South Korea). The primers were tested with PCR using MyTaq DNA polymerase (Bioline, London, UK), with annealing temperature of 61 °C and an extension time of 30 s. Species specificity was tested with DNA from *Ptt*, *Ptm*, *Pyrenophora tritici-repentis, Blumeria graminis* f. sp. *hordei* and barley leaf DNA. The PCR products were visualised by electrophoresis in TAE buffer on a 1.5% agarose gel (Bioline, London, UK) stained with 1x SYBR Safe (Thermo Fisher Scientific, Waltham, MA, USA), run at 90 V for 40 min. Form-specific primers are listed in Table 2.

**Table 2.**
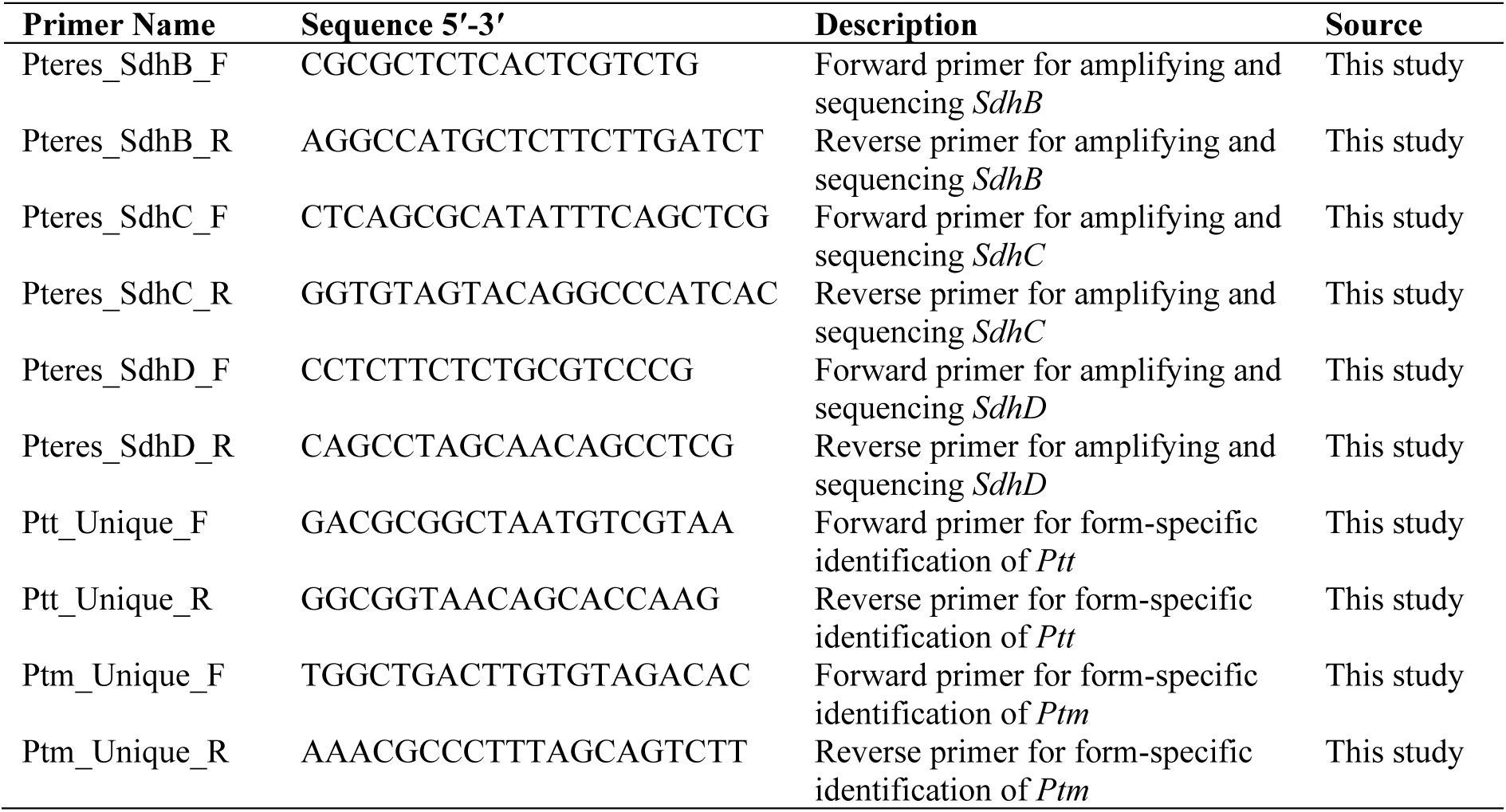
Primers used in this study.

### 2.3 Genotyping of fungicide resistance mutations

Homogenisation of mycelia was conducted with the Mixer Mill MM400 (Retsch GmbH, Haan, Germany), and the extraction of DNA performed in the KingFisher mL Purification System (Thermo Fisher Scientific, Waltham, MA, USA) using the BioSprint 15 DNA Plant Kit (QIAGEN, Hilden, Germany), all according to manufacturers’ instructions. Extracted DNA was quantitated and assessed for purity on a Nanodrop 2000 Spectrophotometer (Thermo Fisher Scientific, Waltham, MA, USA).

PCR amplification of subunits *SdhB*, *SdhC* and *SdhD* was performed with primers listed in Table 2 and the following reaction conditions: PCR amplification used MyTaq DNA Polymerase (Bioline, London, UK), with 0.25 μM each forward and reverse primer, an annealing temperature of 63 °C and an extension time of 1 min. PCR products visualized by electrophoresis on 2% agarose (Bioline, London, UK) gel stained with SYBR Safe (Thermo Fisher Scientific, Waltham, MA, USA) under ultraviolet transillumination. Sanger sequencing was performed on the Applied Biosystems ABI3730XL DNA Analyzer 96-Capillary Array (Thermo Fisher Scientific, Waltham, MA, USA) by Macrogen (Seoul, South Korea). Sequences were aligned using ClustalW algorithm (Thompson *et al*., 1994) with the IUB scoring matrix, gap opening penalty 15, gap extension penalty 6.66 and free end gaps, implemented within Geneious 6.1.8 (Biomatters, Auckland, New Zealand).

### 2.4 *In vitro* fungicide sensitivity phenotyping

Pure technical grade preparations of the SDHI fungicides bixafen, boscalid, fluxapyroxad, isopyrazam and penthiopyrad were used for all assays, original stocks were prepared in ethanol solvent to a concentration of 10 mg mL^-1^. Yeast-Bacto-Acetate liquid medium (10 g yeast extract, 10 g Bacto peptone, 10 g sodium acetate in 1 L sterile deionized H_2_O; YBA) was used for the preparation of 96-well microtiter plate (Corning, NY, USA) assays, as previously described (Mair *et al*., 2016b).

*Ptt* was cultured on V8-Potato-Dextrose Agar (10 g potato-dextrose agar, 3 g CaCO_3_, 15 g agar, 150 mL V8 juice in 850 mL deionized H_2_O; V8PDA) and spores prepared as previously described (Mair *et al*., 2016b). *Ptm* was cultured on Barley Leaf Agar (70 g fresh homogenised barley leaves, 20 g agar in 1 L deionised H_2_O; BLA) and spores prepared as previously described (Mair *et al*., 2020).

A Synergy HT microplate reader (BioTek Instruments, Winooski, VT, USA) with previously described parameters (Mair et al., 2016b) was used for measurement of optical density. Calculation of 50% effective concentrations (EC_50_) was performed as described previously (Mair *et al*., 2016b); log_10_-transformed EC_50_ values (Liang *et al*., 2015) were statistically analysed using Kruskal-Wallis H test and Dunnett’s T3 at the 0.05 significance level, implemented within SPSS Statistics 28 (IBM, Armonk, NY, USA). EC_50_ data visualised with JMP version 16.0 (SAS Institute, Cary, NC, USA). Resistance factors (RF) were calculated as a ratio of the isolate EC_50_ to the mean EC_50_ of 16 wildtype South Australian isolates collected 2015–2019 for *Ptt*; and to the mean EC_50_ of eight wildtype Western Australian isolates collected 1996–2020 for *Ptm*.

### 2.5 Model building and docking

Individual SDH domains were modelled using RoseTTAFold and aligned to the PDB 2WQY. Docking was performed using the software gnina. heme b was flexibly docked on to wildtype, D-H134Y and B-H277L mutants using a flexible D:134 residue and an exhaustiveness of 16, with the autobox ligand binding grid defined from heme b superimposed from PDB 2WQY, all other parameters were set to default settings. Four pyrazole carboxamide fungicides and ubiquinone-2 were flexibly docked onto the above proteins, with heme b bound in the highest scoring modelling pose, using flexible B:233, B:234, B:277, C:63, C:69, C:76, D:46 residues. Exhaustiveness of 16 was used and the autobox ligand binding grid defined from carboxin superimposed from PDB 2WQY, all other parameters were set to default settings.

## 3. Results

### 3.1 Species identification

The form-specific primers successfully amplified only the DNA from the corresponding formae. Electrophoresis of the PCR products with each set of primers revealed bands of the correct size (213bp and 201bp for *Ptt* and *Ptm*, respectively) and no amplification of the closely related fungus *P. tritici-repentis*, the fungus *Blumeria graminis* f. sp. *hordei* which also infects barley or host plant (barley leaf) DNA (figure X). Subsequent testing showed that the *Ptt* differential set also amplifies *Pyrenophora teres* isolated from barley grass (data not shown).

### 3.2 Genotyping of fungicide target genes

#### 3.2.1 South Australian *Ptt*

The *SdhB*, *SdhC* and *SdhD* subunits were sequenced in 36 *Ptt* isolates collected in South Australia in the years 2015 to 2019. All nine of the *Ptt* isolates collected prior to the 2019 season were wildtypes, containing no polymorphisms. In 2019, seven of the sequenced isolates contained no polymorphisms. Of the remaining 20 isolates sequenced from the 2019 season, twelve isolates, collected from seven sites, had a histidine to arginine substitution at amino acid position 134 of the C subunit (C-H134R, GenBank accession number ON534230), which has previously been reported in SDHI resistant *P. teres* in Europe (Rehfus *et al*., 2016). Two isolates from separate sites had an aspartic acid to glycine substitution at amino acid 145 in the D subunit (D-D145G, GenBank accession number ON534237), also seen in SDHI resistant European *P. teres* (Rehfus *et al*., 2016). One isolate had a serine to arginine substitution at amino acid 135 of subunit C via a c489a substitution in the coding sequence (C-S135R, GenBank accession number ON534231), which has been previously observed in SDHI-resistant *P. teres* isolates from Europe (Rehfus *et al*., 2016).

Five isolates from one site in the vicinity of Lock in the Eyre Peninsula had a substitution of histidine to tyrosine at position 134 of subunit D (D-H134Y), this mutation has not been previously described in *P. teres*, although a substitution to arginine (D-H134R) at this position has been identified in SDHI-resistant *P. teres* in Europe (Rehfus *et al*., 2016). The same substitution of histidine to tyrosine has been reported at the homologous position (D-H116Y) in SDHI-resistant *Rhizoctonia cerealis* (Sun *et al*., 2017). The nucleotide sequence of the novel D-H134Y allele has been uploaded to NCBI GenBank (accession number ON534240). No cases were identified of isolates with more than one mutation across the three *Sdh* subunits. A summary of *Ptt* mutations associated with fungicide resistance detected in this study is given in Table 3. Locations where South Australian *Ptt* isolates were collected is summarized in Figure 2.

**Figure 1.**
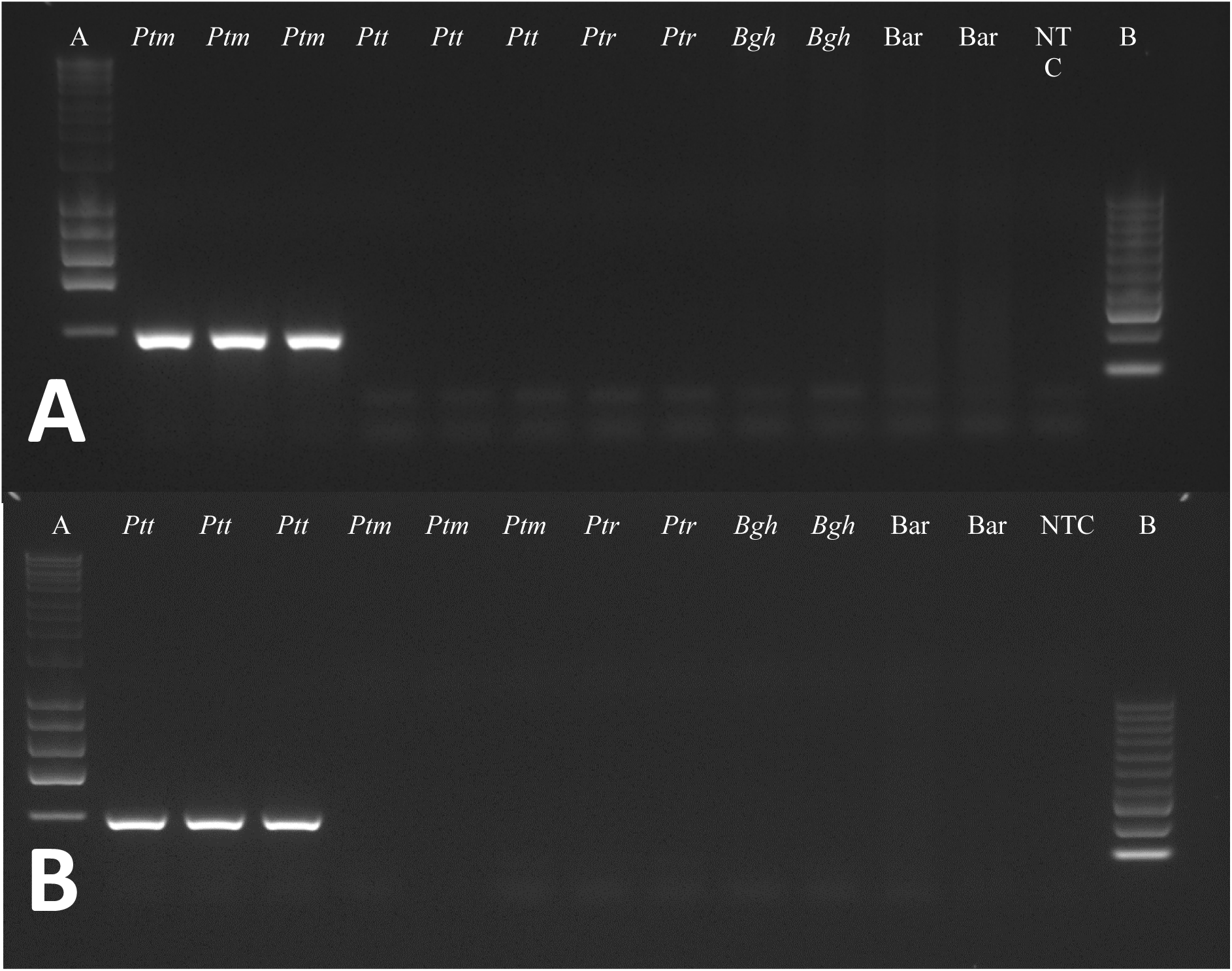
*Ptt* and *Ptm* form-specific PCR markers. **A)** *Ptm* specific primers with DNA from *Ptm* pure culture (wells 2, 3, 4); *Ptt* pure culture (wells 5, 6, 7); *Pyrenophora tritici-repentis* (*Ptr*) pure culture (wells 8, 9); *Blumeria graminis* f.sp. *hordei* (Bgh) DNA (wells 10, 11); Barley DNA (wells 12, 13) and NTC (well 15). Hyperladder 1kb – A (well 1); Hyperladder 100bp – B (well 15) – expected product size 201 bp. **B)** *Ptt* specific primers with DNA from *Ptt* pure culture (wells 2, 3, 4); *Ptm* pure culture (wells 5, 6, 7); *Pyrenophora tritici-repentis* (*Ptr*) pure culture (wells 8, 9); *Blumeria graminis* f.sp. *hordei* (Bgh) DNA (wells 10, 11); Barley DNA (wells 12, 13). Hyperladder 1kb – A (well 1); Hyperladder 100bp – B (well 15) – expected product size 213bp.

**Figure 2.**
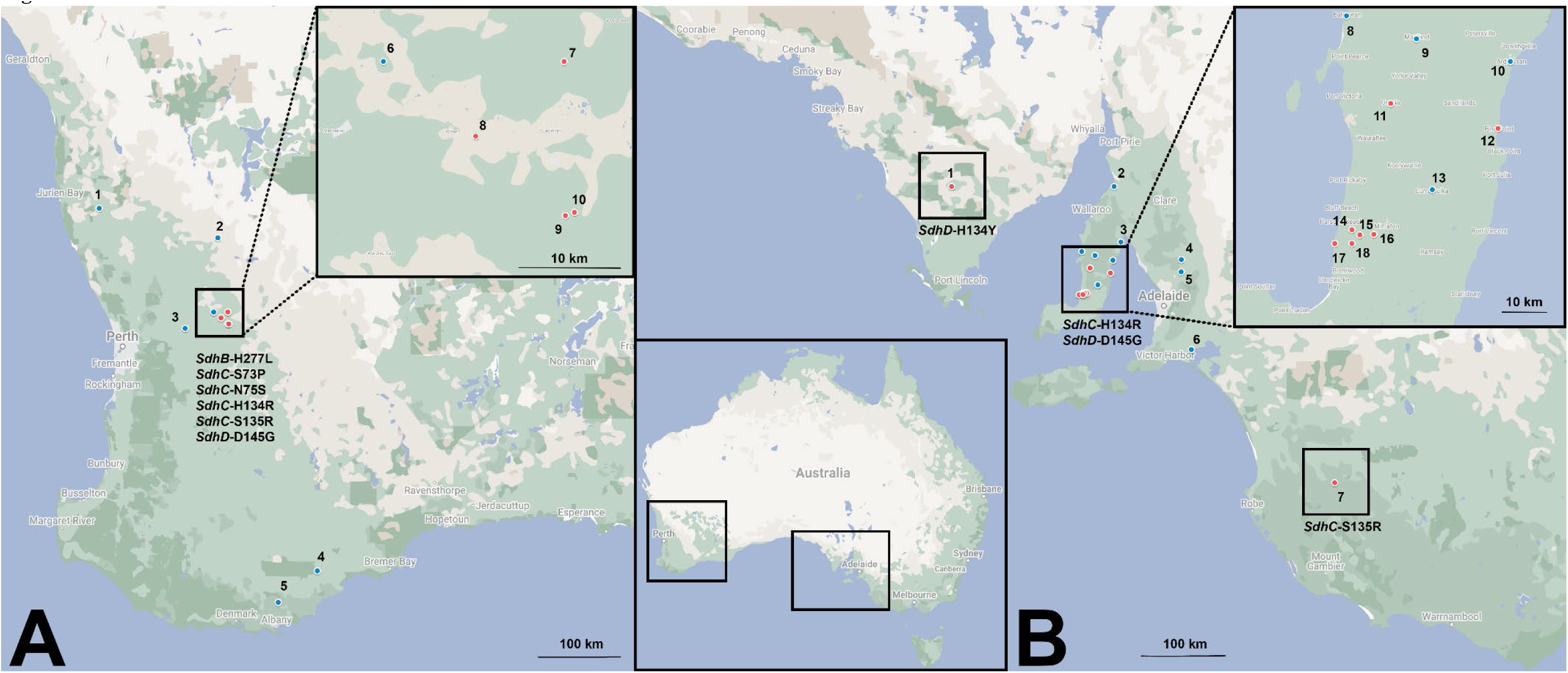
Map of Australia showing geographical origin of *Pyrenophora teres* isolates. **A)** Map of Western Australia showing geographic origin of *Pyrenophora teres* f. sp. *maculata* isolates in this study. Blue: locations where sensitive isolates only were found, red: locations where SDHI-resistant isolates were found. 1. Badgingarra, 2. Cadoux, 3. Muresk, 4. Amelup, 5. Kendenup, 6. Cunderdin (Field 5), 7. Cunderdin (Field 3), 8. Cunderdin (Field 4), 9. Cunderdin (Field 2), 10. Cunderdin (Field 1). **B)** Map of South Australia showing geographic origin of *Pyrenophora teres* f. sp. *teres* isolates in this study. Blue: locations where only sensitive isolates were found, red: locations where SDHI-resistant isolates were found. 1. Lock, 2. Port Broughton, 3. Port Clinton, 4. Freeling, 5. Turretfield, 6. Milang, 7. Kybybolite, 8. Balgowan, 9. Maitland, 10. Ardrossan, 11. Urania, 12. Pine Point, 13. Curramulka, 14. Minlaton (Field 1), 15. Minlaton (Field 2), 16. Minlaton (Field 3), 17. Minlaton (Field 4), 18. Minlaton (Field 5). Map data ©2022 Google.

**Table 3.** Genotypes associated with reduced sensitivity to SDHI fungicides in Australian *Pyrenophora teres* isolates.

| Species | *Sdh*<br>Subunit | Amino Acid<br>Change | Codon WT ><br>Codon Mutant | Isolate(s) | Location | GenBank<br>Accession |
| --- | --- | --- | --- | --- | --- | --- |
| *Ptt* | *SdhC* | H134R | CAC > CGC | 19FRG009, 19FRG011, 19FRG019, 19FRG020,<br>19FRG074, 19FRG077, 19FRG078, 19FRG081,<br>19FRG083, 19FRG085, 19FRG086, 19FRG087 | Yorke Peninsula, SA | ON534230 |
| *Ptt* | *SdhC* | S135R | AGC > AGA | 19FRG091 | Kybybolite, SA | ON534231 |
| *Ptt* | *SdhD* | H134Y | CAC > TAC | 19FRG228, 19FRG229, 19FRG231, 19FRG232,<br>19FRG233 | Eyre Peninsula, SA | ON534240 |
| *Ptt* | *SdhD* | D145G | GAT > GGT | 19FRG010, 19FRG076 | Yorke Peninsula, SA | ON534237 |
| *Ptm* | *SdhB* | H277L | CAC > CTC | 20FRG073 | Cunderdin, WA | ON534228 |
| *Ptm* | *SdhC* | S73P | TCG > CCG | 20FRG041, 20FRG042 | Cunderdin, WA | ON534234 |
| *Ptm* | *SdhC* | N75S | AAC > AGC | 20FRG003, 20FRG004, 20FRG007, 20FRG009,<br>20FRG012 | Cunderdin, WA | ON534233 |
| *Ptm* | *SdhC* | H134R | CAC > CGC | 20FRG011, 20FRG069 | Cunderdin, WA | ON665755 |
| *Ptm* | *SdhC* | S135R | AGC > AGG | 20FRG058 | Cunderdin, WA | ON534235 |
| *Ptm* | *SdhD* | D145G | GAT > GGT | 20FRG001, 20FRG057, 20FRG063, 20FRG068 | Cunderdin, WA | ON665756 |

Species: *Ptt*=*Pyrenophora teres* f. sp. *teres*, *Ptm*=*Pyrenophora teres* f. sp. *maculata*. Location: WA= Western Australia, SA= South Australia.

#### 3.2.2 Western Australian *Ptm*

The *SdhB*, *SdhC* and *SdhD* subunits were sequenced in 17 *Ptm* isolates from Cunderdin, WA collected in the 2020 growing season, as well as in six historical Western Australian *Ptm* isolates. No polymorphisms were observed in the isolates collected prior to 2020. Of the isolates collected in 2020, two isolates were wildtypes, one isolate had the C-S135R substitution via a c489g substitution in the coding sequence (GenBank accession number ON534235), which has also been previously observed in SDHI-resistant *P. teres* isolates from Europe, two isolates had the C-H134R substitution (GenBank accession number ON665755), and four isolates had the D-D145G substitution (GenBank accession number ON665756). Two other isolates had a serine to proline substitution at position 73 of the C subunit (C-S73P, GenBank accession number ON534234), which has been previously observed in SDHI-resistant *P. teres* in Europe (Rehfus *et al*., 2018). One isolate had a histidine to leucine substitution at amino acid 277 of subunit B (B-H277L), this mutation has not been previously described in *P. teres*, although a substitution to tyrosine (B-H277Y) at this position has been identified in SDHI-resistant *P. teres* in Europe (Rehfus *et al*., 2016). The same substitution of histidine to leucine has been reported at the homologous position in SDHI-resistant strains of several species including *Aspergillus oryzae* (Shima *et al*., 2009), *Botrytis cinerea* (FRAC, 2015), *Pyricularia oryzae* (Guo *et al*., 2016), *Ustilago maydis* (Broomfield & Hargreaves, 1992), and *Zymoseptoria tritici* (FRAC, 2015). The nucleotide sequence of the novel B-H277L allele has been uploaded to NCBI GenBank (accession number ON534228). The remaining five *Ptm* isolates had a asparagine to serine substitution at position 75 of the C subunit (C-N75S, GenBank accession number ON534233), which has also previously been identified in SDHI-resistant European populations of *P. teres* (Rehfus *et al*., 2016). As for *Ptt*, no cases were identified of *Ptm* isolates with more than one mutation across the three *Sdh* subunits. A summary of *Ptm* mutations associated with fungicide resistance detected in this study is given in Table 3.

### 3.3 *In vitro* fungicide sensitivity phenotypes

#### 3.3.1 South Australian *Ptt*

36 *Ptt* isolates collected in the years 2015, 2016, 2017 and 2019 from South Australia were tested for sensitivity to the SDHI fungicides boscalid, bixafen, fluxapyroxad, penthiopyrad and isopyrazam (Table 4, Figure 3A). The sixteen isolates with no polymorphisms in *SdhB*, *SdhC* or *SdhD* were classified as wild-type. EC_50_ of sensitive isolates were all significantly (*P* < .001) lower than in isolates with mutations C-H134R, C-S135R, D-H134Y, or D-D145G, and ranged from 0.018 to 0.082 µg mL^-1^ (*M*= 0.041 µg mL^-1^) for boscalid, 0.0019 to 0.0177 µg mL^-1^ (*M*= 0.0065 µg mL^-1^) for bixafen, 0.006 to 0.018 µg mL^-1^ (*M*= 0.010 µg mL^-1^) for fluxapyroxad, 0.013 to 0.110 µg mL^-1^ (*M*= 0.039 µg mL^-1^) for penthiopyrad, and 0.014 to 0.074 µg mL^-1^ (*M*= 0.029 µg mL^-1^) for isopyrazam.

**Figure 3.**
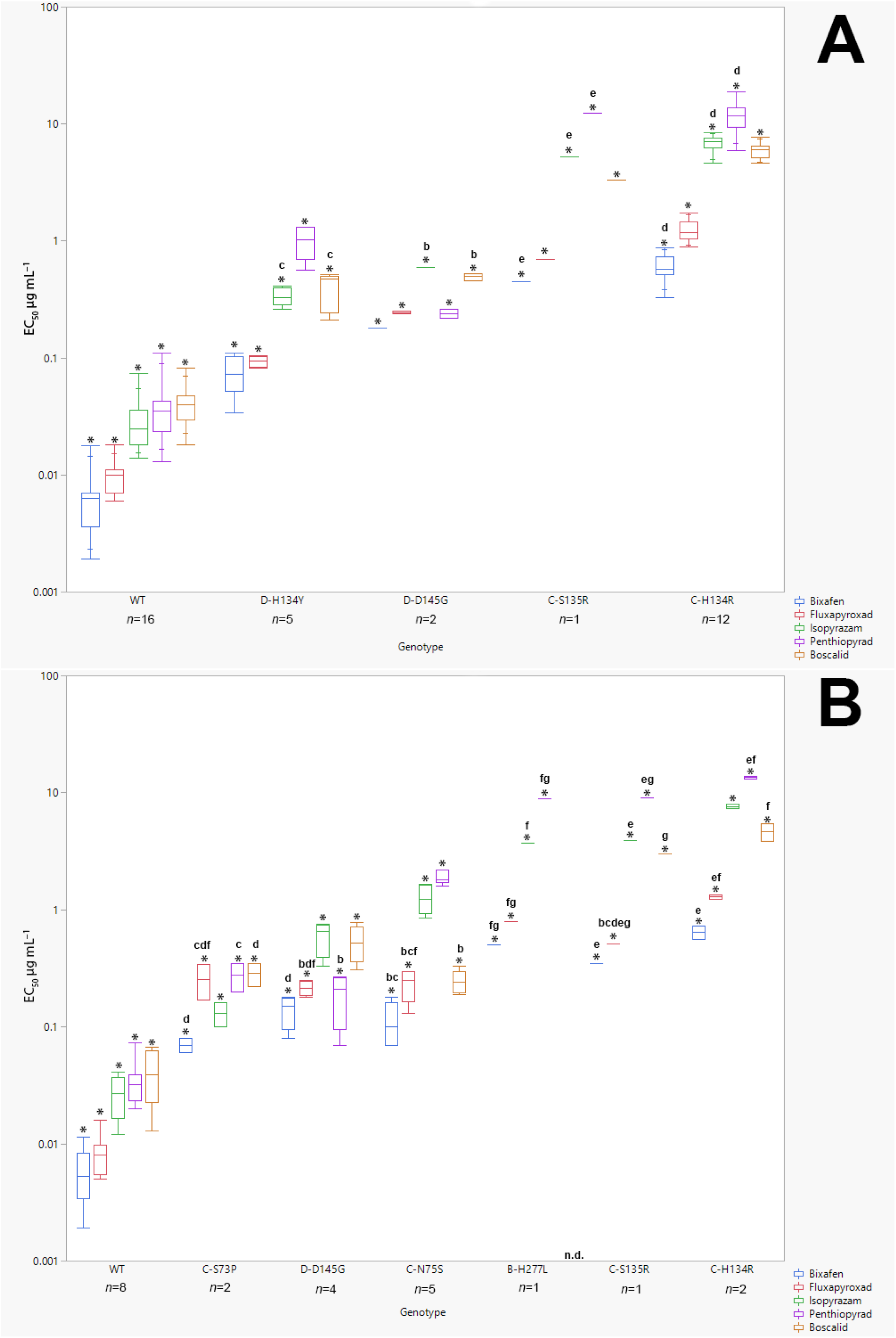
*In vitro* sensitivity of *P yrenophora teres* genotypes to five SDHI fungicides. **A)** *In vitro* sensitivity of *Ptt* genotypes to five SDHI fungicides. Quantile box plots of EC_50_ values (µg mL^-1^) of South Australian *Ptt* isolates. *The mean difference between genotypes within treatments is significant at the .05 level (Dunnett’s T3). Means with same letter within treatments are not significantly different at the .05 level: ^a^WT, ^b^D-H134Y, ^c^D-D145G, ^d^C-S135R, ^e^C-H134R. **B)** In vitro sensitivity of *Ptm* genotypes to five SDHI fungicides. Quantile box plots of EC_50_ values (µg mL^-1^) of Western Australian *Ptm* isolates.*The mean difference between genotypes within treatments is significant at the .05 level (Dunnett’s T3). Means with same letter within treatments are not significantly different at the .05 level: ^a^WT, ^b^C-S73P, ^c^D-D145G, ^d^C-N75S, ^e^B-H277L, ^f^C-S135R, ^g^C-H134R. n.d.: not determined.

**Table 4.** 50% effective concentrations (µg mL^−1^) of *Pyrenophora teres* f. sp. *teres* isolates to five SDHI fungicides.

| Isolate | Boscalid |  | Bixafen |  | Fluxapyroxad |  | Penthiopyrad |  | Isopyrazam |  |
| --- | --- | --- | --- | --- | --- | --- | --- | --- | --- | --- |
|  | EC<sub>50</sub><sup>a</sup> | RF<sup>b</sup> | EC<sub>50</sub> | RF | EC<sub>50</sub> | RF | EC<sub>50</sub> | RF | EC<sub>50</sub> | RF |
| 15FRG252 | 0.043 ($\pm 0.010$) | | 0.0064 ($\pm 0.0010$) | | 0.010 ($\pm 0.001$) | | 0.036 ($\pm 0.004$) | | 0.047 ($\pm 0.009$) | |
| 15FRG253 | 0.029 ($\pm 0.005$) | | 0.0070 ($\pm 0.0012$) | | 0.007 ($\pm 0.001$) | | 0.050 ($\pm 0.007$) | | 0.039 ($\pm 0.012$) | |
| 16FRG031 | 0.044 ($\pm 0.006$) | | 0.0109 ($\pm 0.0026$) | | 0.010 ($\pm 0.001$) | | 0.023 ($\pm 0.004$) | | 0.021 ($\pm 0.003$) | |
| 17FRG007 | 0.040 ($\pm 0.006$) | | 0.0070 ($\pm 0.0009$) | | 0.007 ($\pm 0.001$) | | 0.041 ($\pm 0.007$) | | 0.030 ($\pm 0.002$) | |
| 17FRG021 | 0.042 ($\pm 0.007$) | | 0.0062 ($\pm 0.0005$) | | 0.008 ($\pm 0.001$) | | 0.018 ($\pm 0.005$) | | 0.022 ($\pm 0.004$) | |
| 17FRG022 | 0.051 ($\pm 0.011$) | | 0.0070 ($\pm 0.0006$) | | 0.011 ($\pm 0.002$) | | 0.039 ($\pm 0.011$) | | 0.014 ($\pm 0.003$) | |
| 17FRG052 | 0.026 ($\pm 0.003$) | | 0.0019 ($\pm 0.0005$) | | 0.006 ($\pm 0.001$) | | 0.018 ($\pm 0.002$) | | 0.016 ($\pm 0.003$) | |
| 17FRG111 | 0.049 ($\pm 0.006$) | | 0.0130 ($\pm 0.0025$) | | 0.010 ($\pm 0.002$) | | 0.081 ($\pm 0.020$) | | 0.037 ($\pm 0.006$) | |
| 17FRG112 | 0.082 ($\pm 0.009$) | | 0.0177 ($\pm 0.0028$) | | 0.014 ($\pm 0.003$) | | 0.110 ($\pm 0.013$) | | 0.074 ($\pm 0.010$) | |
| 19FRG046 | 0.065 ($\pm 0.005$) | | 0.0041 ($\pm 0.0011$) | | 0.006 ($\pm 0.001$) | | 0.037 ($\pm 0.010$) | | 0.032 ($\pm 0.006$) | |
| 19FRG056 | 0.032 ($\pm 0.004$) | | 0.0032 ($\pm 0.0013$) | | 0.018 ($\pm 0.004$) | | 0.034 ($\pm 0.005$) | | 0.024 ($\pm 0.005$) | |
| 19FRG075 | 0.034 ($\pm 0.003$) | | 0.0040 ($\pm 0.0003$) | | 0.010 ($\pm 0.001$) | | 0.043 ($\pm 0.003$) | | 0.033 ($\pm 0.003$) | |
| 19FRG080 | 0.035 ($\pm 0.007$) | | 0.0039 ($\pm 0.0006$) | | 0.011 ($\pm 0.003$) | | 0.027 ($\pm 0.004$) | | 0.017 ($\pm 0.004$) | |
| 19FRG082 | 0.018 ($\pm 0.005$) | | 0.0025 ($\pm 0.0008$) | | 0.007 ($\pm 0.001$) | | 0.026 ($\pm 0.009$) | | 0.016 ($\pm 0.004$) | |
| 19FRG084 | 0.040 ($\pm 0.009$) | | 0.0064 ($\pm 0.0016$) | | 0.012 ($\pm 0.001$) | | 0.028 ($\pm 0.006$) | | 0.026 ($\pm 0.005$) | |
| 19FRG088 | 0.025 ($\pm 0.005$) | | 0.0035 ($\pm 0.0008$) | | 0.006 ($\pm 0.002$) | | 0.013 ($\pm 0.004$) | | 0.024 ($\pm 0.006$) | |
| **M=** | **0.041** |  | **0.0065** |  | **0.010** |  | **0.039** |  | **0.029** |  |
| 19FRG228 | 0.28 ($\pm 0.079$) | 6.7 | 0.069 ($\pm 0.014$) | 10.6 | 0.085 ($\pm 0.018$) | 8.8 | 0.56 ($\pm 0.12$) | 14.4 | 0.33 ($\pm 0.06$) | 11.2 |
| 19FRG229 | 0.47 ($\pm 0.037$) | 11.5 | 0.096 ($\pm 0.025$) | 14.7 | 0.082 ($\pm 0.010$) | 8.6 | 1.28 ($\pm 0.06$) | 32.8 | 0.41 ($\pm 0.07$) | 13.8 |
| 19FRG231 | 0.21 ($\pm 0.026$) | 5.1 | 0.034 ($\pm 0.008$) | 5.2 | 0.104 ($\pm 0.013$) | 10.8 | 0.84 ($\pm 0.13$) | 21.4 | 0.31 ($\pm 0.08$) | 10.5 |
| 19FRG232 | 0.52 ($\pm 0.074$) | 12.7 | 0.111 ($\pm 0.023$) | 17.0 | 0.101 ($\pm 0.013$) | 10.5 | 1.03 ($\pm 0.18$) | 26.4 | 0.39 ($\pm 0.05$) | 13.1 |
| 19FRG233 | 0.47 ($\pm 0.088$) | 11.6 | 0.072 ($\pm 0.008$) | 11.0 | 0.094 ($\pm 0.008$) | 9.8 | 1.32 ($\pm 0.07$) | 33.8 | 0.33 ($\pm 0.08$) | 11.2 |
| **M=** | **0.39** | **9.5** | **0.076** | **11.7** | **0.093** | **9.7** | **1.01** | **25.8** | **0.35** | **12.0** |
| 19FRG010 | 0.46 ($\pm 0.02$) | 11.4 | 0.18 ($\pm 0.03$) | 28.1 | 0.24 ($\pm 0.05$) | 25.0 | 0.26 ($\pm 0.02$) | 6.7 | 0.59 ($\pm 0.23$) | 20.0 |
| 19FRG076 | 0.53 ($\pm 0.07$) | 12.9 | 0.18 ($\pm 0.02$) | 26.9 | 0.25 ($\pm 0.04$) | 26.5 | 0.22 ($\pm 0.04$) | 5.6 | 0.89 ($\pm 0.33$) | 30.1 |
| **M=** | **0.50** | **12.1** | **0.18** | **27.5** | **0.25** | **25.8** | **0.24** | **6.2** | **0.74** | **25.0** |
| 19FRG009 | 6.21 ($\pm 0.44$) | 151.9 | 0.59 ($\pm 0.09$) | 90.0 | 1.54 ($\pm 0.27$) | 160.9 | 9.95 ($\pm 1.86$) | 254.5 | 7.66 ($\pm 1.48$) | 260.7 |
| 19FRG011 | 7.66 ($\pm 0.85$) | 187.4 | 0.69 ($\pm 0.15$) | 105.2 | 1.09 ($\pm 0.11$) | 113.1 | 12.13 ($\pm 2.46$) | 310.4 | 7.33 ($\pm 1.48$) | 249.3 |
| 19FRG019 | 5.71 ($\pm 0.35$) | 139.8 | 0.59 ($\pm 0.08$) | 91.1 | 1.42 ($\pm 0.11$) | 148.1 | 11.73 ($\pm 2.05$) | 300.1 | 7.52 ($\pm 1.30$) | 255.8 |
| 19FRG020 | 5.86 ($\pm 0.29$) | 143.4 | 0.53 ($\pm 0.06$) | 81.0 | 1.08 ($\pm 0.10$) | 112.2 | 12.91 ($\pm 0.88$) | 330.3 | 5.78 ($\pm 0.33$) | 196.6 |
| 19FRG074 | 4.64 ($\pm 0.60$) | 113.6 | 0.33 ($\pm 0.05$) | 51.2 | 1.18 ($\pm 0.46$) | 123.4 | 5.90 ($\pm 1.33$) | 150.9 | 7.68 ($\pm 1.71$) | 261.4 |
| 19FRG077 | 6.42 ($\pm 0.64$) | 157.2 | 0.50 ($\pm 0.11$) | 76.7 | 0.89 ($\pm 0.14$) | 92.9 | 11.21 ($\pm 2.10$) | 286.8 | 4.67 ($\pm 0.71$) | 159.0 |
| 19FRG078 | 5.16 ($\pm 0.47$) | 126.3 | 0.52 ($\pm 0.11$) | 79.9 | 1.46 ($\pm 0.31$) | 152.4 | 9.10 ($\pm 1.71$) | 232.7 | 7.01 ($\pm 0.94$) | 238.6 |
| 19FRG081 | 6.76 ($\pm 0.41$) | 165.4 | 0.87 ($\pm 0.13$) | 132.9 | 1.75 ($\pm 0.18$) | 182.8 | 18.83 ($\pm 2.34$) | 481.6 | 8.44 ($\pm 1.17$) | 287.3 |
| 19FRG083 | 6.43 ($\pm 0.36$) | 157.4 | 0.79 ($\pm 0.15$) | 120.5 | 1.16 ($\pm 0.17$) | 121.2 | 14.04 ($\pm 3.49$) | 359.3 | 7.02 ($\pm 0.77$) | 238.9 |
| 19FRG085 | 5.16 (±0.52) | 126.3 | 0.55 (±0.09) | 84.3 | 0.99 (±0.12) | 103.3 | 18.92 (±0.95) | 484.1 | 5.95 (±0.57) | 202.5 |
| 19FRG086 | 4.90 (±0.90) | 120.0 | 0.52 (±0.17) | 79.7 | 1.44 (±0.29) | 149.7 | 9.13 (±2.19) | 233.6 | 6.96 (±1.39) | 236.9 |
| 19FRG087 | 6.18 (±0.67) | 151.2 | 0.75 (±0.26) | 115.3 | 1.03 (±0.22) | 107.7 | 11.76 (±1.85) | 300.8 | 7.01 (±0.99) | 238.5 |
| **M=** | **5.92** | **145.0** | **0.60** | **92.3** | **1.25** | **130.7** | **12.13** | **310.6** | **6.92** | **235.5** |
| 19FRG091 | 3.33 (±0.24) | 81.5 | 0.45 (±0.05) | 68.7 | 0.70 (±0.06) | 72.8 | 12.37 (±1.19) | 316.5 | 5.23 (±0.51) | 178.0 |

<sup>a</sup>EC<sub>50</sub> (50% effective concentration) values are the mean of at least three biological replicates (± Standard Error). <sup>b</sup>Resistance Factor: ratio of the isolate EC<sub>50</sub> to the mean EC<sub>50</sub> of sensitive isolates.

EC_50_ values were significantly (*P* < .005) higher in isolates with the C-H134R mutation compared with all other genotypes for boscalid (4.64-7.66 µg mL^-1^, *M*= 5.92 µg mL^-1^) and fluxapyroxad (0.89-1.75 µg mL^-1^, *M*= 1.25 µg mL^-1^). EC_50_ values of C-H134R isolates for bixafen (0.33-0.87 µg mL^-1^, *M*= 0.60 µg mL^-1^), penthiopyrad (5.9-18.9 µg mL^-1^, *M*= 12.1 µg mL^-1^) and isopyrazam (4.67-8.44 µg mL^-1^, *M*= 6.92 µg mL^-1^) were not significantly different (*P* > .05) from those of the isolate with the mutation C-S135R (*M*= 0.45, 12.4 and 5.23 µg mL^-1^, respectively).

EC_50_ values of the isolate with the C-S135R mutation were significantly (*P* < .001) higher than those with D-H134Y or D-D145G for all fungicides, and second-highest overall for boscalid (3.33 µg mL^-1^) and fluxapyroxad (0.70 µg mL^-1^).

EC_50_ of the isolate with the mutation D-D145G was significantly (*P* < .001) higher than those with D-H134Y for both bixafen (mean EC_50_ 0.18 and 0.076 µg mL^-1^, respectively) and fluxapyroxad (*M*= 0.25 and 0.093 µg mL^-1^, respectively), and significantly (*P* < .001) lower for penthiopyrad (*M*= 0.24 and 1.0 µg mL^-1^, respectively).

Differences between EC_50_ of isolates with the mutations D-H134Y and D-D145G were not significant for boscalid (*P* = .099) or isopyrazam (*P* = .886).

#### 3.3.2 Western Australian *Ptm*

23 *Ptm* isolates collected between 1996 and 2020 from Western Australian were tested for sensitivity to the SDHI fungicides boscalid, bixafen, fluxapyroxad, penthiopyrad and isopyrazam (Table 5, Figure 3B). The eight isolates with no polymorphisms in *SdhB*, *SdhC* or *SdhD* were classified as wild-type for the purpose of calculating Resistance Factors. EC_50_ of sensitive isolates were all significantly (*P* < .001) lower than in isolates with any mutation, and ranged from 0.013 to 0.067 µg mL^-1^ (*M*= 0.040 µg mL^-1^) for boscalid, 0.0019 to 0.0115 µg mL^-1^ (*M*= 0.0050 µg mL^-1^) for bixafen, 0.002 to 0.016 µg mL^-1^ (*M*= 0.008 µg mL^-1^) for fluxapyroxad, 0.023 to 0.073 µg mL^-1^ (*M*= 0.038 µg mL^-1^) for penthiopyrad, and 0.014 to 0.041 µg mL^-1^ (*M*= 0.030 µg mL^-1^) for isopyrazam.

**Table 5.**
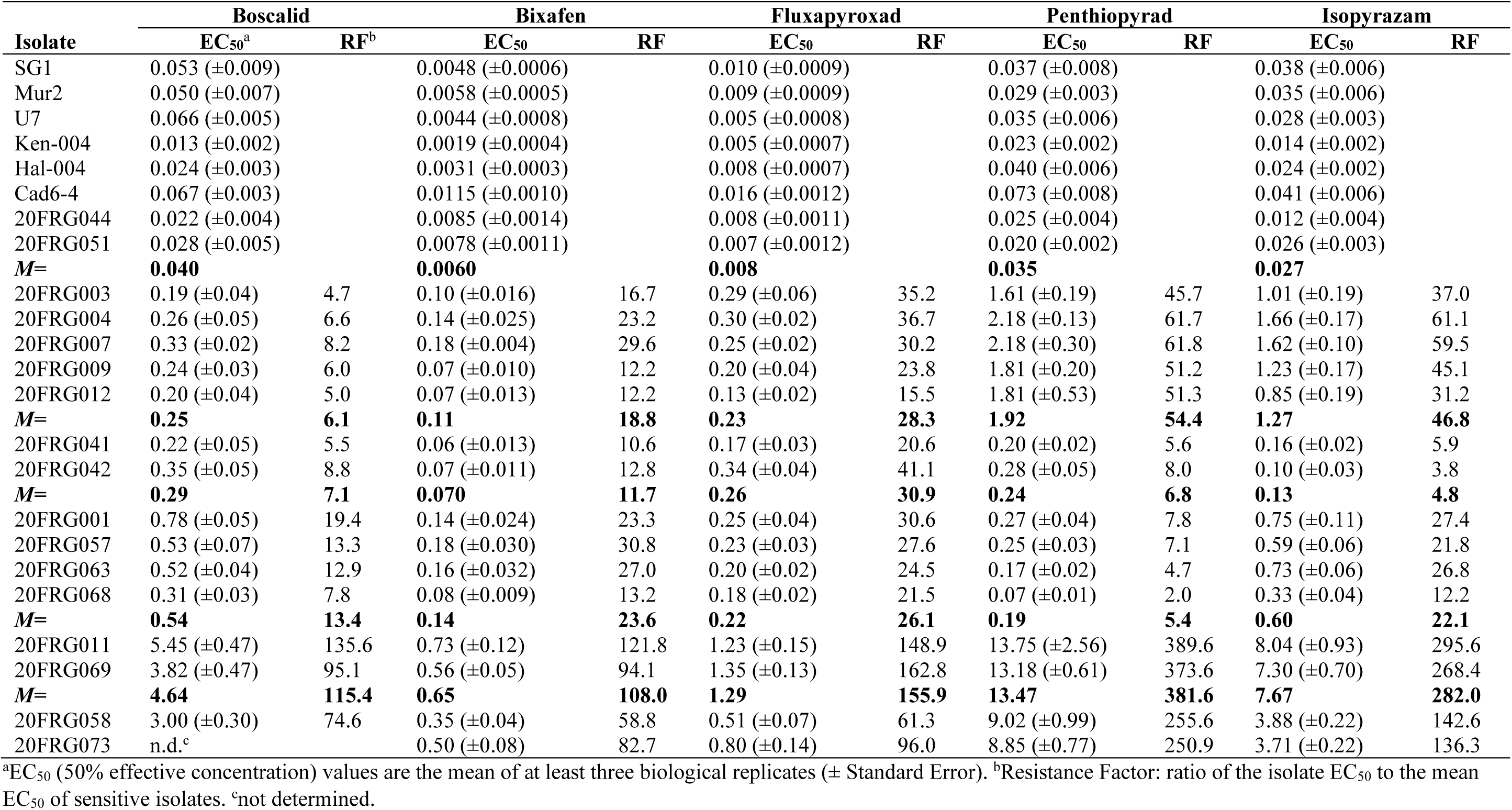
50% effective concentrations (µg mL^−1^) of *Pyrenophora teres* f. sp. *maculata* isolates to five SDHI fungicides.

| Isolate | Boscalid |  | Bixafen |  | Fluxapyroxad |  | Penthiopyrad |  | Isopyrazam |  |
| --- | --- | --- | --- | --- | --- | --- | --- | --- | --- | --- |
|  | EC<sub>50</sub><sup>a</sup> | RF<sup>b</sup> | EC<sub>50</sub> | RF | EC<sub>50</sub> | RF | EC<sub>50</sub> | RF | EC<sub>50</sub> | RF |
| SG1 | 0.053 ($\pm 0.009$) | | 0.0048 ($\pm 0.0006$) | | 0.010 ($\pm 0.0009$) | | 0.037 ($\pm 0.008$) | | 0.038 ($\pm 0.006$) | |
| Mur2 | 0.050 ($\pm 0.007$) | | 0.0058 ($\pm 0.0005$) | | 0.009 ($\pm 0.0009$) | | 0.029 ($\pm 0.003$) | | 0.035 ($\pm 0.006$) | |
| U7 | 0.066 ($\pm 0.005$) | | 0.0044 ($\pm 0.0008$) | | 0.005 ($\pm 0.0008$) | | 0.035 ($\pm 0.006$) | | 0.028 ($\pm 0.003$) | |
| Ken-004 | 0.013 ($\pm 0.002$) | | 0.0019 ($\pm 0.0004$) | | 0.005 ($\pm 0.0007$) | | 0.023 ($\pm 0.002$) | | 0.014 ($\pm 0.002$) | |
| Hal-004 | 0.024 ($\pm 0.003$) | | 0.0031 ($\pm 0.0003$) | | 0.008 ($\pm 0.0007$) | | 0.040 ($\pm 0.006$) | | 0.024 ($\pm 0.002$) | |
| Cad6-4 | 0.067 ($\pm 0.003$) | | 0.0115 ($\pm 0.0010$) | | 0.016 ($\pm 0.0012$) | | 0.073 ($\pm 0.008$) | | 0.041 ($\pm 0.006$) | |
| 20FRG044 | 0.022 ($\pm 0.004$) | | 0.0085 ($\pm 0.0014$) | | 0.008 ($\pm 0.0011$) | | 0.025 ($\pm 0.004$) | | 0.012 ($\pm 0.004$) | |
| 20FRG051 | 0.028 ($\pm 0.005$) | | 0.0078 ($\pm 0.0011$) | | 0.007 ($\pm 0.0012$) | | 0.020 ($\pm 0.002$) | | 0.026 ($\pm 0.003$) | |
| **M=** | **0.040** |  | **0.0060** |  | **0.008** |  | **0.035** |  | **0.027** |  |
| 20FRG003 | 0.19 ($\pm 0.04$) | 4.7 | 0.10 ($\pm 0.016$) | 16.7 | 0.29 ($\pm 0.06$) | 35.2 | 1.61 ($\pm 0.19$) | 45.7 | 1.01 ($\pm 0.19$) | 37.0 |
| 20FRG004 | 0.26 ($\pm 0.05$) | 6.6 | 0.14 ($\pm 0.025$) | 23.2 | 0.30 ($\pm 0.02$) | 36.7 | 2.18 ($\pm 0.13$) | 61.7 | 1.66 ($\pm 0.17$) | 61.1 |
| 20FRG007 | 0.33 ($\pm 0.02$) | 8.2 | 0.18 ($\pm 0.004$) | 29.6 | 0.25 ($\pm 0.02$) | 30.2 | 2.18 ($\pm 0.30$) | 61.8 | 1.62 ($\pm 0.10$) | 59.5 |
| 20FRG009 | 0.24 ($\pm 0.03$) | 6.0 | 0.07 ($\pm 0.010$) | 12.2 | 0.20 ($\pm 0.04$) | 23.8 | 1.81 ($\pm 0.20$) | 51.2 | 1.23 ($\pm 0.17$) | 45.1 |
| 20FRG012 | 0.20 ($\pm 0.04$) | 5.0 | 0.07 ($\pm 0.013$) | 12.2 | 0.13 ($\pm 0.02$) | 15.5 | 1.81 ($\pm 0.53$) | 51.3 | 0.85 ($\pm 0.19$) | 31.2 |
| **M=** | **0.25** | **6.1** | **0.11** | **18.8** | **0.23** | **28.3** | **1.92** | **54.4** | **1.27** | **46.8** |
| 20FRG041 | 0.22 ($\pm 0.05$) | 5.5 | 0.06 ($\pm 0.013$) | 10.6 | 0.17 ($\pm 0.03$) | 20.6 | 0.20 ($\pm 0.02$) | 5.6 | 0.16 ($\pm 0.02$) | 5.9 |
| 20FRG042 | 0.35 ($\pm 0.05$) | 8.8 | 0.07 ($\pm 0.011$) | 12.8 | 0.34 ($\pm 0.04$) | 41.1 | 0.28 ($\pm 0.05$) | 8.0 | 0.10 ($\pm 0.03$) | 3.8 |
| **M=** | **0.29** | **7.1** | **0.070** | **11.7** | **0.26** | **30.9** | **0.24** | **6.8** | **0.13** | **4.8** |
| 20FRG001 | 0.78 ($\pm 0.05$) | 19.4 | 0.14 ($\pm 0.024$) | 23.3 | 0.25 ($\pm 0.04$) | 30.6 | 0.27 ($\pm 0.04$) | 7.8 | 0.75 ($\pm 0.11$) | 27.4 |
| 20FRG057 | 0.53 ($\pm 0.07$) | 13.3 | 0.18 ($\pm 0.030$) | 30.8 | 0.23 ($\pm 0.03$) | 27.6 | 0.25 ($\pm 0.03$) | 7.1 | 0.59 ($\pm 0.06$) | 21.8 |
| 20FRG063 | 0.52 ($\pm 0.04$) | 12.9 | 0.16 ($\pm 0.032$) | 27.0 | 0.20 ($\pm 0.02$) | 24.5 | 0.17 ($\pm 0.02$) | 4.7 | 0.73 ($\pm 0.06$) | 26.8 |
| 20FRG068 | 0.31 ($\pm 0.03$) | 7.8 | 0.08 ($\pm 0.009$) | 13.2 | 0.18 ($\pm 0.02$) | 21.5 | 0.07 ($\pm 0.01$) | 2.0 | 0.33 ($\pm 0.04$) | 12.2 |
| **M=** | **0.54** | **13.4** | **0.14** | **23.6** | **0.22** | **26.1** | **0.19** | **5.4** | **0.60** | **22.1** |
| 20FRG011 | 5.45 ($\pm 0.47$) | 135.6 | 0.73 ($\pm 0.12$) | 121.8 | 1.23 ($\pm 0.15$) | 148.9 | 13.75 ($\pm 2.56$) | 389.6 | 8.04 ($\pm 0.93$) | 295.6 |
| 20FRG069 | 3.82 ($\pm 0.47$) | 95.1 | 0.56 ($\pm 0.05$) | 94.1 | 1.35 ($\pm 0.13$) | 162.8 | 13.18 ($\pm 0.61$) | 373.6 | 7.30 ($\pm 0.70$) | 268.4 |
| **M=** | **4.64** | **115.4** | **0.65** | **108.0** | **1.29** | **155.9** | **13.47** | **381.6** | **7.67** | **282.0** |
| 20FRG058 | 3.00 ($\pm 0.30$) | 74.6 | 0.35 ($\pm 0.04$) | 58.8 | 0.51 ($\pm 0.07$) | 61.3 | 9.02 ($\pm 0.99$) | 255.6 | 3.88 ($\pm 0.22$) | 142.6 |
| 20FRG073 | n.d.<sup>c</sup> | | 0.50 ($\pm 0.08$) | 82.7 | 0.80 ($\pm 0.14$) | 96.0 | 8.85 ($\pm 0.77$) | 250.9 | 3.71 ($\pm 0.22$) | 136.3 |

<sup>a</sup>EC<sub>50</sub> (50% effective concentration) values are the mean of at least three biological replicates ( ± Standard Error). <sup>b</sup>Resistance Factor: ratio of the isolate EC<sub>50</sub> to the mean EC<sub>50</sub> of sensitive isolates. <sup>c</sup>not determined.

Isolates with the mutations B-H277L, C-H134R, or C-S135R had significantly (*P* < .05) higher EC_50_ values for bixafen, penthiopyrad, and isopyrazam, compared to those with mutations *SdhC*-S73P, *SdhC*-N75S, or *SdhD*-D145G. EC_50_ values for isopyrazam were significantly (*P* < .001) higher in the isolates with the C-H134R mutation (*M*= 7.7 µg mL^-1^) compared with all other genotypes. Differences between EC_50_ of isolates with the mutations C-H134R and C-S135R were not significant for boscalid (*M*= 4.6 and 3.0 µg mL^-1^, respectively; *P* = .392), penthiopyrad (*M*= 13.5 and 9.0 µg mL^-1^, respectively; *P* = .179), or fluxapyroxad (*M*= 1.29 and 0.51 µg mL^-1^, respectively; *P* = .177). Differences between EC_50_ of isolates with the mutations C-H134R and B-H277L were not significant for bixafen (*M*= 0.65 and 0.50 µg mL^-1^, respectively; *P* = .995), penthiopyrad (*M*= 13.5 and 8.9 µg mL^-1^, respectively; *P* = .170), or fluxapyroxad (*M*= 1.29 and 0.80 µg mL^-1^, respectively; *P* = .503). Differences between EC_50_ of isolates with the mutations C-S135R and B-H277L were not significant for bixafen (*M*= 0.35 and 0.50 µg mL^-1^, respectively; *P* = .938), penthiopyrad (*M*= 9.0 and 8.9 µg mL^-1^, respectively; *P* = 1), fluxapyroxad (*M*= 0.51 and 0.80 µg mL^-1^, respectively; *P* = .869), or isopyrazam (*M*= 3.88 and 3.71 µg mL^-1^, respectively; *P* = 1), sensitivity of isolate with the mutation B-H277L to boscalid was not determined.

Differences between mean EC_50_ of isolates with the C-S73P, D-D145G mutation and C-N75S mutations were significant (*P* < .001) for isopyrazam (*M*= 0.13, 0.60, and 1.27 µg mL^-1^, respectively). EC_50_ values were significantly higher (*P* < .001) for penthiopyrad in isolates with C-N75S (*M*= 1.92 µg mL^-1^) compared to those with C-S73P (*M*= 0.24 µg mL^-1^) or D-D145G (*M*= 0.19 µg mL^-1^), difference between C-S73P and D-D145G for penthiopyrad were not significant (*P* = .917). EC_50_ values were significantly higher (*P* ≤ .002) for boscalid in isolates with D-D145G (*M*= 0.54 µg mL^-1^) compared to those with C-S73P (*M*= 0.29 µg mL^-1^) or C-N75S (*M*= 0.25 µg mL^-1^), difference between C-S73P and D-D145G for penthiopyrad were not significant (*P* = .906). The EC_50_ to bixafen for isolates with the C-N75S mutation (*M*= 0.11 µg mL^-1^) was not significantly different to those with C-S73P (*M*= 0.07 µg mL^-1^, P= .270) or D-D145G (*M*= 0.14 µg mL^-1^, P= .792). The differences between EC_50_ values to fluxapyroxad were not significant (P > .05) for isolates with C-S73P (*M*= 0.26 µg mL^-1^), C-N75S (*M*= 0.23 µg mL^-1^), D-D145G (*M*= 0.22 µg mL^-1^) or C-S135R (*M*= 0.51 µg mL^-1^).

#### 3.3.3 Comparison of genotypes in *Ptt* and *Ptm*

There was no significant differences between the EC_50_ values of the 16 wild-type *Ptt* isolates and the eight wild-type *Ptm* isolates for any of the five SDHI fungicides (*P* > .05). There was no significant differences between the EC_50_ values of the *Ptt* and *Ptm* isolates carrying the C-H134R, C-S135R, or D-D145G mutations, respectively, for any of the five SDHI fungicides (*P* > .05).

### 3.4 Modelling reveals molecular basis for fungicide resistance in D-H134Y and B-H277L mutants

Prediction of the *P. teres* SDH structure using the software RoseTTAFold (Baek *et al*., 2021) revealed a highly similar fold to previous SDH crystal structures and placed the D-H134Y mutation within the heme b binding regions which would likely affect heme b binding (Figure 4). Docking of the heme into this region revealed a different binding pose to wildtype likely the result of steric hindrance from the larger tyrosine sidechain. Interestingly, the heme b was shifted towards the ubiquinone binding site (Q-site)(Sun *et al*., 2005; Baek *et al*., 2021) where it could also affect inhibitor binding and local to B-H277. Docking of four pyrazole carboxamide fungicides into this region in wildtype *P. teres* SDH revealed a similar binding mode (Sup. Fig. 1). Docking of penthiopyrad, the pyrazole carboxamide fungicide to which most resistance was seen, to D-H134Y and B-H277L mutants revealed highly altered binding modes. Notably, penthiopyrad was not found in the ubiquinone binding pocket for the highest scoring modelling pose in B-H277L mutant which also gave the greatest resistance *in vivo* (Figure 5). Therefore, the increased resistance seen in this mutant, in comparison to the D-H134Y mutation, is likely due to the direct effects of the hydrophobic leucine sidechain distorting the canonical hydrophilic interactions between H277 and the thiophene and carbonyl groups of the pyrazole carboxamide fungicides. Whereas the reduced resistance in the D-H134Y mutation is likely due to the indirect effect of an altered heme b binding mode hindering fungicide binding. Notably, ubiquinone was predicted to bind in a similar manner to the ubiquinone binding site in wildtype and mutants illustrating how *P. teres* SDH may still function in the presence of these mutation (Sup. Fig. 2).

**Figure 4.**
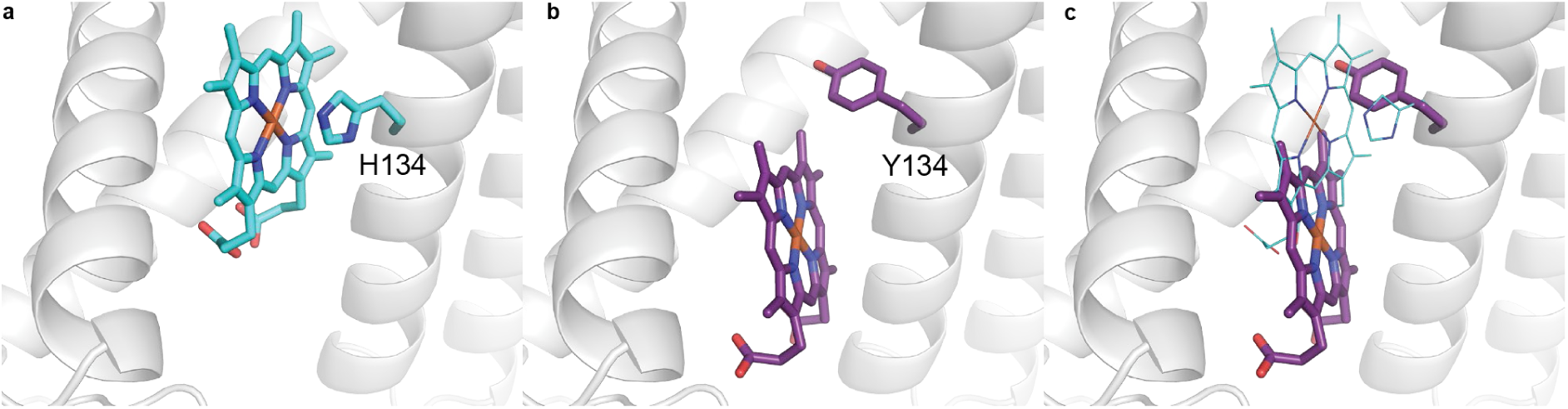
Modelling predicts heme *b* binding to be altered in D-H134Y mutant. A model of *P. teres* SDH was made using the RoseTTAFold (Ref.1) and heme *b* docked into the SDH-D domain (grey cartoon) using gnina (Ref.2). (**a**) In the highest scoring modelling pose heme *b* (cyan sticks) is docked adjacent to flexible wildtype SDH-D-His134 residue (labelled cyan sticks). (**b**) Mutation of His134 to Tyr134 (labelled purple sticks) results in an altered binding mode of heme *b* (purple sticks) likely induced by steric hindrance from Tyr134. (**c**) An overlay of **a** and **b** illustrating altered heme *b* binding between Tyr134 mutant (purple sticks) and wildtype (cyan lines).

**Figure 5.**
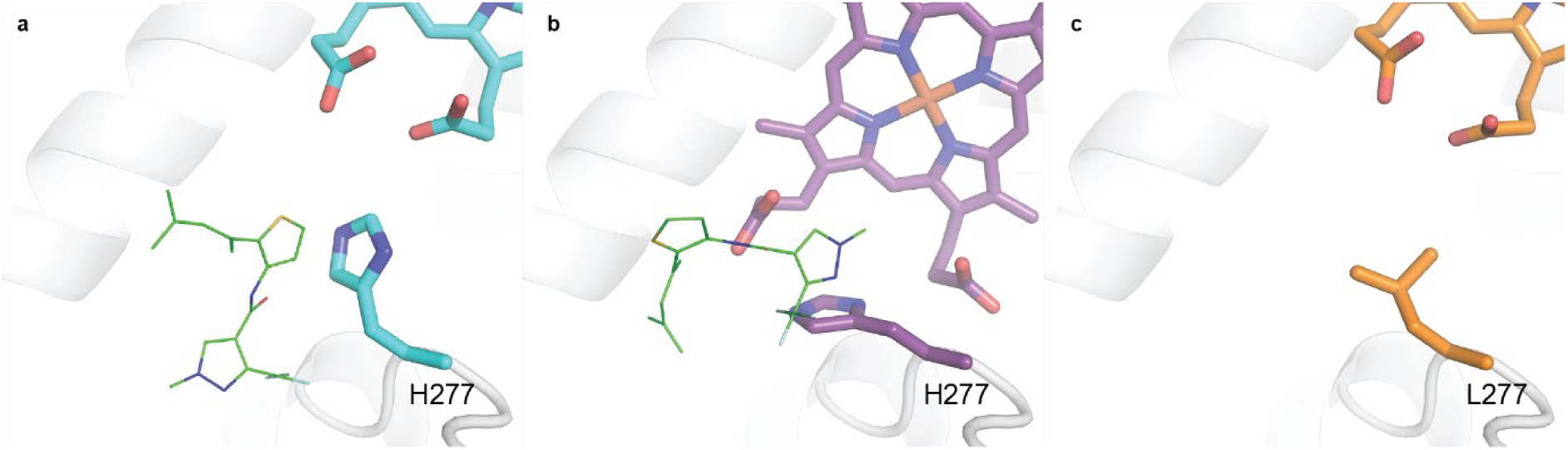
Modelling predicts penthiopyrad binding to be altered in D-H134Y and B-H277L mutants. Penthiopyrad (green lines) was docked into heme *b* containing *P **yrenophora** teres* SDH wildtype (a) and D-H134Y (b) and B-H277L (c) mutants using gnina. (**a**) In the highest scoring modelling pose the thiophene and carbonyl group of penthiopyrad are adjacent to wildtype SDH B-H277 in the ubiquinone binding site. (**b**) Penthiopyrad binds in the same region to wildtype in the D-H134Y mutant but in a different conformation due to the shifted heme *b*. (**c**) The presence of the more hydrophobic B-H277L mutant resulted in penthiopyrad binding outside of the wildtype ubiquinone binding site in the highest scoring modelling pose.

## 4. Discussion

The first cases of SDHI fungicide resistance in *P. teres* were observed in 2019 in South Australia, and in Western Australia the following year. Therefore, the emergence of resistance to SDHIs occurred within three years of the first introduction of these compounds, indicating a particularly rapid development of resistance. In Europe, SDHI resistance in *P. teres* emerged six and five years after the introduction of these compounds to Germany and France, respectively (Rehfus, 2018). *P. teres* is polycyclic, with up to 12 infection cycles per year depending on conditions (Rehfus, 2018); and is considered a medium-risk pathogen for fungicide resistance development (FRAC, 2019). Higher disease pressure and a greater number of infection cycles can contribute to the more rapid development of resistance (Rehfus, 2018). Therefore, it is possible that more conducive conditions in Australia have contributed to the more rapid emergence of resistance. An alternative explanation is the incursion of resistant genotypes into Australia from Europe or other sources. However, the occurrence of unique resistance mutations (B-H277L, D-H134Y) not previously observed in other continents may militate against this hypothesis. The emergence of the same amino acid substitution (C-S135R) in South Australian *Ptt* and Western Australian *Ptm* populations via two different nucleotide changes in the coding sequence (c489a versus c489g) also supports the possibility of repeated, independent parallel evolution of these traits, as opposed to a model of gene flow where resistance is gained by the spread of mutant alleles from a point of origin; however both point mutations have been observed in C-S135R *P. teres* isolates from Europe (Rehfus *et al*., 2016). Hartmann *et al*. (2021), in an analysis of *Z. tritici* populations from Oregon, Switzerland and Australia, found that *Cyp51* resistance mutations likely emerged rapidly and independently in North American and European populations, while there was also evidence of gene flow of resistant alleles from Europe to Australia. Future population genetic or population genomic studies of South Australian, Western Australia, European and other populations of *P. teres* may help elucidate the origins of SDHI-resistance mutations and the contribution of *de novo* mutation versus gene flow.

Over 27 mutations in the Sdh complex have so far been observed across phytopathogenic fungi (Shi *et al*., 2021). The most frequently mutated residue is at *H277* of the B subunit (position number based on archetype *SdhB* sequence from *P. teres*, (Mair *et al*., 2016c)), which has been found to be mutated in every species for which SDHI resistance has been observed (Sierotzki & Scalliet, 2013a), most frequently substituted to a tyrosine at this position (Lalève *et al*., 2014). In *P. teres*, the C-*G79R* mutation has previously been reported to be the most common, representing over half of all resistant European isolates, with B-*H277Y* the next most common mutation (Rehfus, 2018). However, neither mutation was observed in Australian isolates in this study: there were no mutations observed at residue 79 of *SdhC* in any isolate, while a single isolate had a histidine to leucine substitution at residue 277 of *SdhB* (B-*H277L*), which has not previously been described in *P. teres*. The B-*H277Y* mutation in *P. teres* was reported by Rehfus (2018) as being associated with among the lowest resistance factors to all the tested SDHI compounds, whereas B-*H277L* in this study is associated with among the highest resistance factors. These observations are congruent with those previously reported for other pathogens such as *Botrytis cinerea* (Leroux *et al*., 2010; Veloukas *et al*., 2013), *Magnaporthe oryzae* (Liu *et al*., 2022) and *Zymoseptoria tritici* (Scalliet *et al*., 2012), where a leucine substitution is associated with much higher resistance factors than a tyrosine substitution at the equivalent *H277* position of *SdhB*. A possible explanation then for the comparative rarity of the B-*H277L* mutation is that, although associated with high resistance factors, it has also been associated with the largest reduction in Sdh enzyme activity (Lalève *et al*., 2014). For instance, Lalève *et al*. (2014) found that the B-H272Y mutation in *Botrytis cinerea* (B-*H277Y* in *P. teres*) was the only mutation to have no effect on respiration rate or Sdh activity of the mutations studied, whereas B-H272L (B-*H277L*) was associated with the lowest respiration rate, 58% lower than the wild-type. Conversely, the D-*H134Y* mutation described in this study was associated with lower resistance factors than the D-*H134R* mutation described by Rehfus (2018). As arginine is not a coordination partner for haem b (Dokmanić *et al*., 2008), the D-*H134R* substitution results in a loss of a coordination partner for the haem b in the mutant Sdh complex (Rehfus, 2018), which may potentially result in a decrease in reduction potential of the haem b iron, or loss of the haem b group causing a decrease in structural stability of the enzyme (Stammler *et al*., 2015). In contrast, tyrosine is able to act as a coordination partner of haem (Li *et al*., 2011), though docking of haem b to a Sdh complex with the D-*H134Y* substitution reveals a different binding pose compared to wild-type, caused by steric hindrance from the larger tyrosine sidechain. A potential direction for future research is to investigate the effect of these various substitution on Sdh enzymatic activity compared to the wild-type, with its attendant implications for overall fitness.

The EC_50_ values and resistance factors described in this study are in general higher than those previously described in European *P. teres* (Rehfus *et al*., 2016; Rehfus, 2018; Stammler, 2021), most likely a reflection of differences in experimental method for EC_50_ measurement between the studies, as well as differences between the individual isolates used. As EC_50_ values often vary greatly between laboratories they generally are not directly comparable; rather, it is the rank order of fungicides and genotypes that should be compared (Landschoot *et al*., 2017). And in fact the rank order of resistance factors for the four mutations described in both studies (C-H134R, C-S135R, C-N75S and D-D145G) and the four fungicides tested in both studies (bixafen, fluxapyroxad, isopyrazam and penthiopyrad) are all identical (Table 6).

**Table 6.**
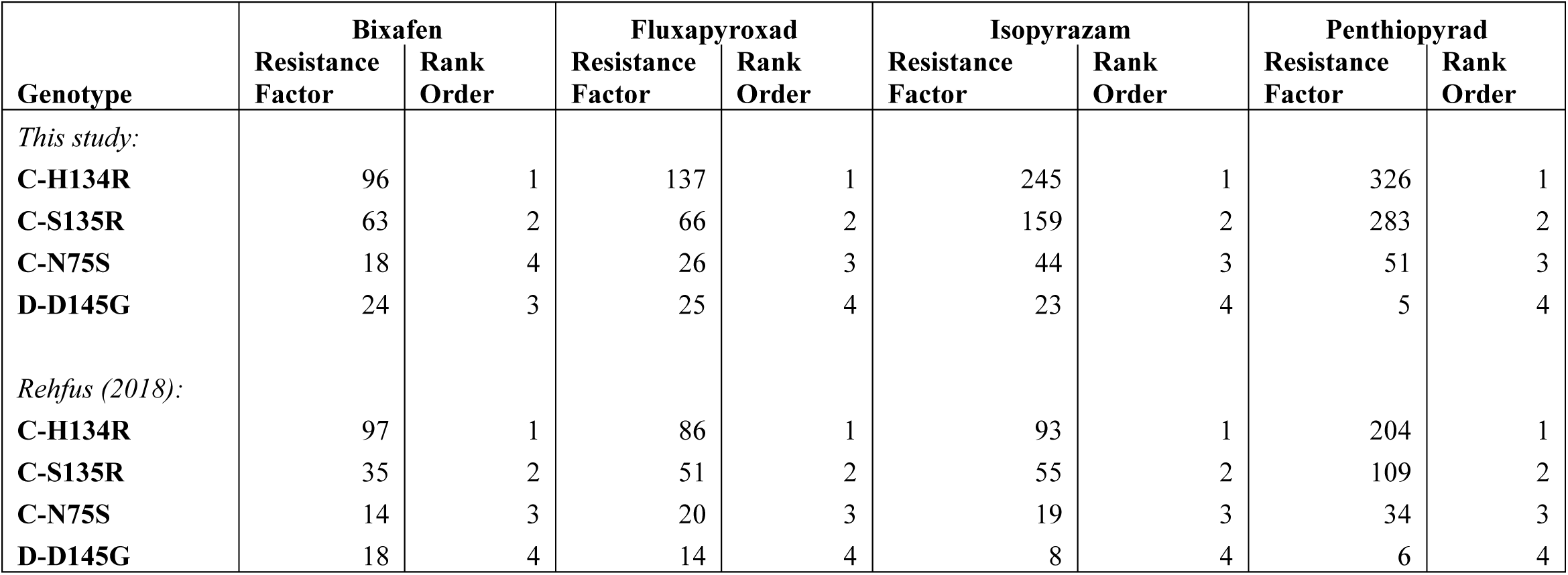
Comparison of mean resistance factors of *Pyrenophora teres* with different mutations in the Sdh complex.

| **Genotype** | **Bixafen** |  | **Fluxapyroxad** |  | **Isopyrazam** |  | **Penthiopyrad** |  |
| --- | --- | --- | --- | --- | --- | --- | --- | --- |
|  | **Resistance Factor** | **Rank Order** | **Resistance Factor** | **Rank Order** | **Resistance Factor** | **Rank Order** | **Resistance Factor** | **Rank Order** |
| *This study:* |  |  |  |  |  |  |  |  |
| **C-H134R** | 96 | 1 | 137 | 1 | 245 | 1 | 326 | 1 |
| **C-S135R** | 63 | 2 | 66 | 2 | 159 | 2 | 283 | 2 |
| **C-N75S** | 18 | 4 | 26 | 3 | 44 | 3 | 51 | 3 |
| **D-D145G** | 24 | 3 | 25 | 4 | 23 | 4 | 5 | 4 |
| *Rehfus (2018):* |  |  |  |  |  |  |  |  |
| **C-H134R** | 97 | 1 | 86 | 1 | 93 | 1 | 204 | 1 |
| **C-S135R** | 35 | 2 | 51 | 2 | 55 | 2 | 109 | 2 |
| **C-N75S** | 14 | 3 | 20 | 3 | 19 | 3 | 34 | 3 |
| **D-D145G** | 18 | 4 | 14 | 4 | 8 | 4 | 6 | 4 |

The emergence of SDHI resistance in both net form and spot form of net blotch represents a challenge for the Australian barley industry, particularly in light of the ongoing issues of DMI resistance in both *Ptt* (Mair *et al*., 2016b) and *Ptm* (Mair *et al*., 2020) in Australia. There is therefore an urgent need for integrated disease management, such as improved crop rotations and stubble management, to reduce disease pressure and prolong the effective lifespan of SDHI fungicides.

## Acknowledgements

The authors would like to thank the sample contributions made by growers and agronomists. Primers for Sdh genes designed by Dr. Katherine Zulak, Curtin University. This study was conducted by the Centre for Crop and Disease Management, a joint initiative of Curtin University and the Grains Research and Development Corporation (research grant CUR00023).

---

**Supplementary Figure 1.**
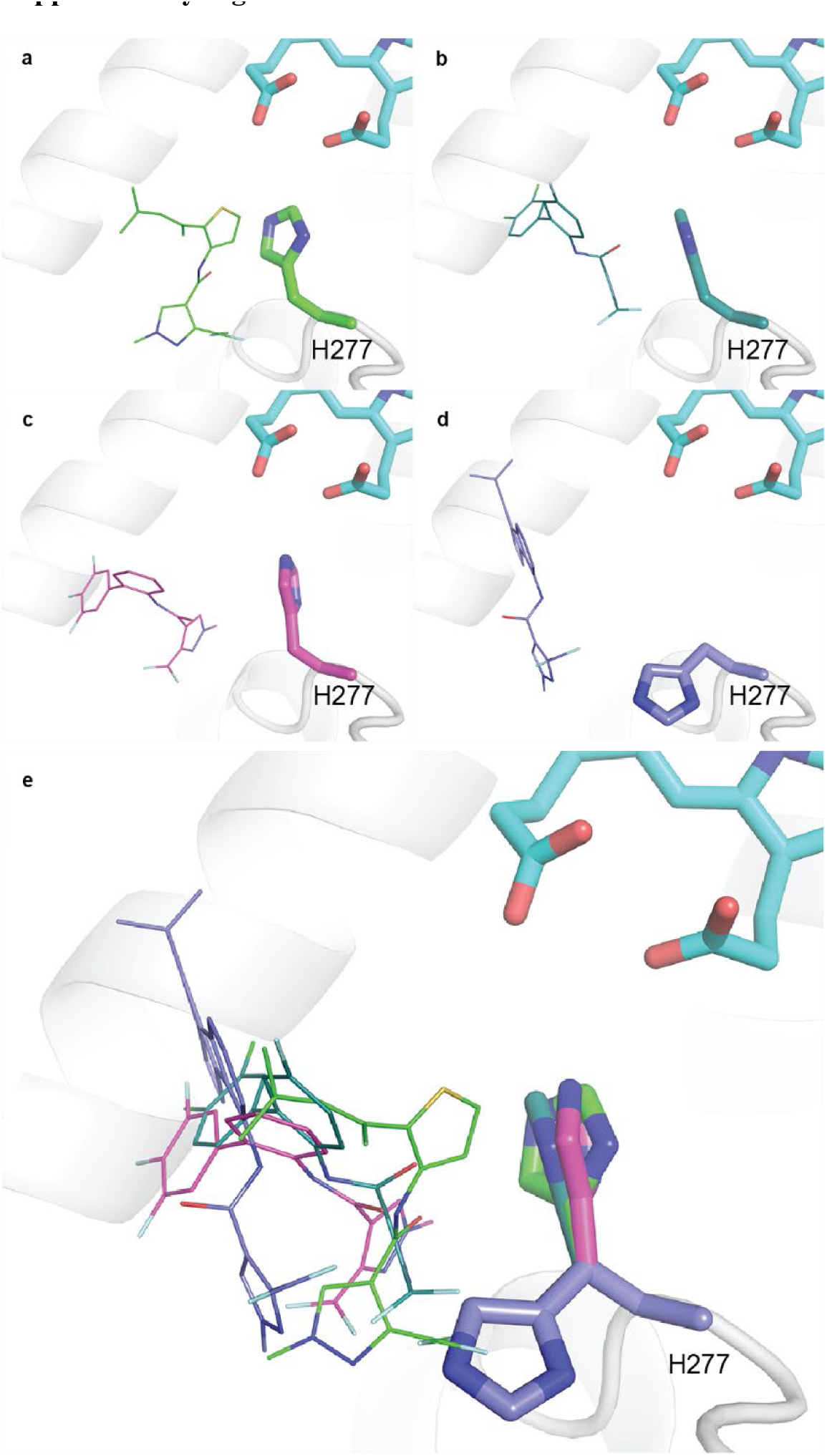
Modelling predicts similar binding modes for four pyrazole carboxamide fungicides in *P yrenophora teres* SDH. Wildtype SDH (grey cartoon) with H277 residue shown as sticks in same colour as bound fungicide in the highest scoring modelling pose. (**a**) Penthiopyrad (green line). (**b**) Bixafen (teal line). (**c**) Fluxapyroxad (magenta line). (**d**) Isopyrazam (blue line). (**e**) Overlay of a-d reveals the flexibility of H277 in pyrazole carboxamide fungicides binding and their similar binding modes with pyrazole domain buried into the ubiquinone binding pocket adjacent to H277.

**Supplementary Figure 2.**
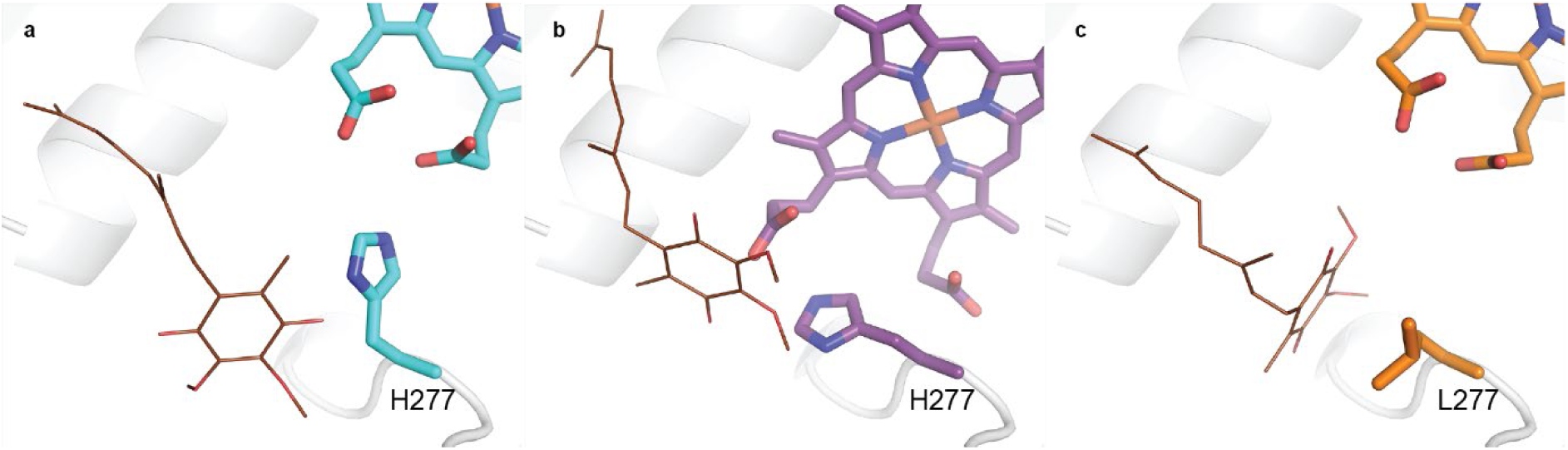
Modelling predicts ubiquinone to bind wildtype, D-H134Y and B-H277L mutants in the ubiquinone binding site. Ubiquinone-2 (brown lines) was docked into heme *b* containing *P yrenophora teres* SDH wildtype (a) and D-H134Y (b) and B-H277L (c) mutants using gnina. (a) In the highest scoring modelling pose the quinone group is found within the ubiquinone binding site local to residue D-277 (labelled).

## References

1. Akhavan A, Strelkov SE, Askarian H, Kher SV, Fraser M, Kutcher HR, Turkington TK. 2017. Sensitivity of western Canadian Pyrenophora teres f. teres and P. teres f. maculata isolates to propiconazole and pyraclostrobin. Canadian Journal of Plant Pathology 39(1): 11–24.

2. APVMA 2022. PubCRIS: Public Chemical Registration Information System Search. Australia: Australian Pesticides and Veterinary Medicines Authority.

3. Avenot HF, Michailides TJ. 2010. Progress in understanding molecular mechanisms and evolution of resistance to succinate dehydrogenase inhibiting (SDHI) fungicides in phytopathogenic fungi. Crop Protection 29(7): 643–651.

4. Baek M, DiMaio F, Anishchenko I, Dauparas J, Ovchinnikov S, Lee GR, Wang J, Cong Q, Kinch LN, Schaeffer RD, et al. 2021. Accurate prediction of protein structures and interactions using a three-track neural network. Science 373(6557): 871–876.

5. Bayer 2018. Global fungicide resistance experts tour Australia: Bayer Australia.

6. Broomfield PLE, Hargreaves JA. 1992. A single amino-acid change in the iron-sulphur protein subunit of succinate dehydrogenase confers resistance to carboxin in Ustilago maydis. Current genetics 22(2): 117–121.

7. Dokmanić I, Sikic M, Tomić S. 2008. Metals in Proteins: Correlation Between the Metal-Ion Type, Coordination Number and the Amino-Acid Residues Involved in the Coordination. Acta crystallographica. Section D, Biological crystallography 64: 257–263.

8. Ellwood SR, Syme RA, Moffat CS, Oliver RP. 2012. Evolution of three Pyrenophora cereal pathogens: Recent divergence, speciation and evolution of non-coding DNA. Fungal Genetics and Biology 49(10): 825–829.

9. FRAC 2015. List of Species Resistant to SDHIs April 2015: Fungicide Resistance Action Committee (FRAC).

10. FRAC 2019. FRAC Pathogen Risk List 2019 Fungicide Resistance Action Committee.

11. Guo M, Zhu X, Li H, Tan L, Pan Y. 2016. Development of a novel strategy for fungal transformation based on a mutant locus conferring carboxin-resistance in Magnaporthe oryzae. AMB Express 6(1): 57.

12. Hartmann FE, Vonlanthen T, Singh NK, McDonald MC, Milgate A, Croll D. 2021. The complex genomic basis of rapid convergent adaptation to pesticides across continents in a fungal plant pathogen. Molecular Ecology 30(21): 5390–5405.

13. Kretschmer M, Leroch M, Mosbach A, Walker A-S, Fillinger S, Mernke D, Schoonbeek H-J, Pradier J-M, Leroux P, De Waard MA, et al. 2009. Fungicide-Driven Evolution and Molecular Basis of Multidrug Resistance in Field Populations of the Grey Mould Fungus Botrytis cinerea. PLOS Pathogens 5(12): e1000696.

14. Lalève A, Gamet S, Walker AS, Debieu D, Toquin V, Fillinger S. 2014. Site-directed mutagenesis of the P225, N230 and H272 residues of succinate dehydrogenase subunit B from Botrytis cinerea highlights different roles in enzyme activity and inhibitor binding. Environmental Microbiology 16(7): 2253–2266.

15. Landschoot S, Carrette J, Vandecasteele M, De Baets B, Höfte M, Audenaert K, Haesaert G. 2017. Boscalid-resistance in Alternaria alternata and Alternaria solani populations: An emerging problem in Europe. Crop Protection 92: 49–59.

16. Leroux P, Gredt M, Leroch M, Walker A-S. 2010. Exploring Mechanisms of Resistance to Respiratory Inhibitors in Field Strains of Botrytis cinerea, the Causal Agent of Gray Mold. Applied and environmental microbiology 76(19): 6615–6630.

17. Li T, Bonkovsky HL, Guo J-t. 2011. Structural analysis of heme proteins: implications for design and prediction. BMC structural biology 11: 13–13.

18. Liang HJ, Li JL, Di YL, Zhang AS, Zhu FX. 2015. Logarithmic transformation is essential for statistical analysis of fungicide EC50 values. Journal of Phytopathology 163(6): 456–464.

19. Liu S, Ma J, Jiang B, Yang G, Guo M. 2022. Functional characterization of MoSdhB in conferring resistance to pydiflumetofen in blast fungus Magnaporthe oryzae. Pest management science 78(10): 4018–4027.

20. Liu Z, Ellwood SR, Oliver RP, Friesen TL. 2011. Pyrenophora teres: profile of an increasingly damaging barley pathogen. Molecular Plant Pathology 12(1): 1–19.

21. Mair WJ, Deng W, Mullins JG, West S, Wang P, Besharat N, Ellwood SR, Oliver RP, Lopez-Ruiz FJ. 2016a. Demethylase Inhibitor Fungicide Resistance in Pyrenophora teres f. sp. teres Associated with Target Site Modification and Inducible Overexpression of Cyp51. Frontiers in microbiology 7: 1279.

22. Mair WJ, Deng W, Mullins JG, West S, Wang P, Besharat N, Ellwood SR, Oliver RP, Lopez-Ruiz FJ. 2016b. Demethylase Inhibitor Fungicide Resistance in Pyrenophora teres f. sp. teres Associated with Target Site Modification and Inducible Overexpression of Cyp51. Front Microbiol 7: 1279.

23. Mair WJ, Lopez-Ruiz F, Stammler G, Clark W, Burnett F, Hollomon D, Ishii H, Thind TS, Brown JK, Fraaije B, et al. 2016c. Proposal for a unified nomenclature for target-site mutations associated with resistance to fungicides. Pest management science 72(8): 1449–1459.

24. Mair WJ, Thomas GJ, Dodhia K, Hills AL, Jayasena KW, Ellwood SR, Oliver RP, Lopez-Ruiz FJ. 2020. Parallel evolution of multiple mechanisms for demethylase inhibitor fungicide resistance in the barley pathogen Pyrenophora teres f. sp. maculata. Fungal Genetics and Biology 145: 103475.

25. Murray GM, Brennan JP. 2010. Estimating disease losses to the Australian barley industry. Australasian Plant Pathology 39(1): 85–96.

26. Omrane S, Sghyer H, Audéon C, Lanen C, Duplaix C, Walker A-S, Fillinger S. 2015. Fungicide efflux and the MgMFS1 transporter contribute to the multidrug resistance phenotype in Zymoseptoria tritici field isolates. Environmental Microbiology 17(8): 2805–2823.

27. Pearce TL, Wilson CR, Gent DH, Scott JB. 2019. Multiple mutations across the succinate dehydrogenase gene complex are associated with boscalid resistance in Didymella tanaceti in pyrethrum. PloS one 14(6): e0218569.

28. Platz G, Snyman L, Fowler R 2017. Systiva performance in northern trials: Grains Research and Development Corporation.

29. Rehfus A. 2018. Analysis of the emerging situation of resistance to succinate dehydrogenase inhibitors in Pyrenophora teres and Zymoseptoria tritici in Europe. Universität Hohenheim.

30. Rehfus A, Miessner S, Achenbach J, Strobel D, Bryson R, Stammler G. 2016. Emergence of succinate dehydrogenase inhibitor resistance of Pyrenophora teres in Europe. Pest management science 72(10): 1977–1988.

31. Rehfus A, Strobel D, Bryson R, Stammler G. 2018. Mutations in sdh genes in field isolates of Zymoseptoria tritici and impact on the sensitivity to various succinate dehydrogenase inhibitors. Plant Pathology 67(1): 175–180.

32. Samaras Α, Ntasiou P, Myresiotis C, Karaoglanidis G. 2020. Multidrug resistance of Penicillium expansum to fungicides: whole transcriptome analysis of MDR strains reveals overexpression of efflux transporter genes. International Journal of Food Microbiology 335: 108896.

33. Sang H, Hulvey J, Popko JT, Jr., Lopes J, Swaminathan A, Chang T, Jung G. 2015. A pleiotropic drug resistance transporter is involved in reduced sensitivity to multiple fungicide classes in Sclerotinia homoeocarpa (F.T. Bennett). Molecular Plant Pathology 16(3): 251–261.

34. Scalliet G, Bowler J, Luksch T, Kirchhofer-Allan L, Steinhauer D, Ward K, Niklaus M, Verras A, Csukai M, Daina A, et al. 2012. Mutagenesis and Functional Studies with Succinate Dehydrogenase Inhibitors in the Wheat Pathogen Mycosphaerella graminicola. PloS one 7(4): e35429.

35. Serenius M, Manninen O. 2008. Prochloraz tolerance of Pyrenophora teres population in Finland. Agricultural and food science 15(1): 35–42.

36. Sheridan J, Grbavac, Sheridan MH, Soteros J. 1987. Further studies on triadimenol resistance in the barley net blotch pathogen Pyrenophora teres. New Zealand Journal of Agricultural Research 30(1): 101–105.

37. Sheridan JE, Grbavac N, Sheridan MH. 1985. Triadimenol insensitivity in Pyrenophora teres. Transactions of the British Mycological Society 85(2): 338–341.

38. Shi Y, Zhu F, Sun B, Xie X, Chai A, Li B. 2021. Two adjacent mutations in the conserved domain of SdhB confer various resistance phenotypes to fluopyram in Corynespora cassiicola. Pest Manag Sci 77(9): 3980–3989.

39. Shima Y, Ito Y, Kaneko S, Hatabayashi H, Watanabe Y, Adachi Y, Yabe K. 2009. Identification of three mutant loci conferring carboxin-resistance and development of a novel transformation system in Aspergillus oryzae. Fungal Genetics and Biology 46(1): 67–76.

40. Sierotzki H, Frey R, Wullschleger J, Palermo S, Karlin S, Godwin J, Gisi U. 2007. Cytochrome b gene sequence and structure of Pyrenophora teres and P. tritici-repentis and implications for QoI resistance. Pest management science 63(3): 225–233.

41. Sierotzki H, Scalliet G. 2013a. A Review of Current Knowledge of Resistance Aspects for the Next-Generation Succinate Dehydrogenase Inhibitor Fungicides. Phytopathology® 103(9): 880–887.

42. Sierotzki H, Scalliet G. 2013b. A review of current knowledge of resistance aspects for the next-generation succinate dehydrogenase inhibitor fungicides. Phytopathology 103(9): 880–887.

43. Stammler G 2021. personal communication: BASF.

44. Stammler G, Wolf A, Glaettli A, Klappach K 2015. Respiration Inhibitors: Complex II. In: Ishii H, Hollomon DW eds. Fungicide Resistance in Plant Pathogens: Principles and a Guide to Practical Management. Tokyo: Springer Japan, 105–117.

45. Steinhauer D, Salat M, Frey R, Mosbach A, Luksch T, Balmer D, Hansen R, Widdison S, Logan G, Dietrich RA, et al. 2019. A dispensable paralog of succinate dehydrogenase subunit C mediates standing resistance towards a subclass of SDHI fungicides in Zymoseptoria tritici. PLOS Pathogens 15(12): e1007780.

46. Sun F, Huo X, Zhai Y, Wang A, Xu J, Su D, Bartlam M, Rao Z. 2005. Crystal Structure of Mitochondrial Respiratory Membrane Protein Complex II. Cell 121(7): 1043–1057.

47. Sun H-Y, Lu C-Q, Li W, Deng Y-Y, Chen H-G. 2017. Homozygous and heterozygous point mutations in succinate dehydrogenase subunits b, c and d of Rhizoctonia cerealis conferring resistance to thifluzamide. Pest management science 73(5): 896–903.

48. Syme RA, Martin A, Wyatt NA, Lawrence JA, Muria-Gonzalez MJ, Friesen TL, Ellwood SR. 2018. Transposable Element Genomic Fissuring in Pyrenophora teres Is Associated With Genome Expansion and Dynamics of Host–Pathogen Genetic Interactions. Frontiers in Genetics 9(130).

49. Thompson JD, Higgins DG, Gibson TJ. 1994. CLUSTAL W: improving the sensitivity of progressive multiple sequence alignment through sequence weighting, position-specific gap penalties and weight matrix choice. Nucleic acids research 22(22): 4673–4680.

50. Veloukas T, Markoglou AN, Karaoglanidis GS. 2013. Differential Effect of SdhB Gene Mutations on the Sensitivity to SDHI Fungicides in Botrytis cinerea. Plant Dis 97(1): 118–122.

51. Yamashita M, Fraaije B. 2018. Non-target site SDHI resistance is present as standing genetic variation in field populations of Zymoseptoria tritici. Pest management science 74(3): 672–681.

52. Zhang Y, Lu J, Wang J, Zhou M, Chen C. 2015. Baseline sensitivity and resistance risk assessmemt of Rhizoctonia cerealis to thifluzamide, a succinate dehydrogenase inhibitor. Pestic Biochem Physiol 124: 97–102.

